# Super-resolved imaging and Bayesian analysis of single exocytosis events reveals molecular-scale patterning by cortical microtubule arrays

**DOI:** 10.1101/2024.12.01.626273

**Authors:** Jelmer J. Lindeboom, Ryan Gutierrez, Viktor Kirik, David W. Ehrhardt

## Abstract

The microtubule cytoskeleton organizes exocytosis to enable cellular morphogenesis, but how non-centrosomal arrays control exocytotic site positioning remains poorly understood. Using elongating plant cells as a model, we developed quantitative methods to move beyond coarse correlation and reveal the precise spatial relationship between cortical microtubules and secretion. We identify KEULE, an essential SEC/MUNC protein, as a dynamic exocytosis marker that forms clusters with stereotyped assembly and disassembly kinetics at discrete secretion sites. Combining confocal microscopy with super-resolution analysis and Bayesian inference, we quantified microtubule-exocytosis positioning at nanometer precision. This analysis revealed that microtubules create ∼180 nm enrichment zones flanked by ∼520 nm depletion zones, generating a spatial pattern that replicates the cortical array structure. Unexpectedly, Bayesian inference showed strong evidence for a flat enrichment profile within these zones rather than peaked distributions. This flat profile, combined with enrichment zone widths exceeding the ∼50 nm reach of known tethering proteins, challenges a direct vesicle capture mechanism. Instead, our data support a membrane domain mechanism where microtubules organize lipid/protein composition to create preferred exocytosis territories. These findings establish quantitative spatial rules for how non-centrosomal microtubule arrays organize secretion.

## Introduction

The ability to organize biomolecules at the cell cortex was a fundamental innovation in the evolution of cellular life, enabling essential biological functions including cellular morphogenesis, motility and polarized transport. Targeted exocytosis, non-random positioning and fusion of exocytotic vesicles at the plasma membrane, is a crucial mechanism for creating such cortical organization. Across phyla, the actin and microtubule cytoskeletons have been implicated in spatial organization of exocytosis. This functionality is exemplified in plant and fungal cells by cellular tip growth, and in animal cells by polarization of mammalian epithelia, cell migration and the positioning of focal adhesions^1,2^. The molecular polarity of microtubules plays an important role in how their function in exocytotic targeting is understood. For example, polarized arrays of microtubules created by nucleation from centralized organizing bodies, such as centrosomes or Golgi, can serve as directional tracks for motor-driven transport of exocytotic vesicles from where they are created in the cell interior to peripheral destinations at the plasma membrane^2,3^. More refined positioning of exocytosis is proposed to be accomplished by association of microtubule plus ends with specific locations at the plasma membrane, including focal adhesions^2^. Polarized arrays of microtubules can also organize exocytosis through directed transport of factors that establish membrane domains that serve to target exocytotic traffic, as during tip growth in fission yeast^4^. However, many cells across diverse organisms feature interphase microtubule arrays that are not centrally organized and have architectures where microtubules are not clearly polarized with respect to the plasma membrane^5^. Non-centrosomal arrays are required in important biological processes that require exocytosis, such as morphogenesis, but relatively little is understood about how they may function to target and pattern exocytotic traffic.

Gaining an understanding of exocytotic targeting mechanisms requires an ability to observe individual exocytotic events with respect to targeting machineries and a means to verify successful delivery of exocytotic cargo. Observing individual exocytosis events has been challenging in many cell models because microtubules, exocytotic vesicles and the machineries of exocytosis are often densely concentrated. Axially growing cells in the epidermis of higher plants have features that provide an excellent experimental platform to make such observations and for investigating how non-centrosomal microtubule arrays may target and pattern exocytosis. In contrast to cells with microtubule arrays that are polarized with respect to the cell interior and the plasma membrane, these cells feature highly ordered arrays of microtubules that lie at the cell cortex in parallel to and in contact with the plasma membrane along their lattices. Large regions of the plasma membrane lie in a single focal plane, where the individual microtubules and microtubule bundles that constitute these arrays are easily visualized due to their quasi-2-dimensional nature and comparatively low density^6^.

In previous studies, we determined that cellulose synthase complexes (CSCs), large integral membrane protein complexes that synthesize and extrude cellulose into the cell wall, can be used to observe individual exocytotic events at the plasma membrane by allowing dynamic tracking of both exocytotic vesicles and the successful delivery of cargo to the plasma membrane^7–9^. By tracking CSCs while simultaneously imaging dynamic cortical arrays, we discovered that CSCs were preferentially delivered to the plasma membrane at locations defined by microtubules. This targeting did not involve vesicular transport along microtubules, nor did it involve defining exocytotic domains by end-on association of microtubule ends with the plasma membrane^7^, thus suggesting a previously undescribed mechanism for microtubule-dependent exocytotic targeting. However, CSC’s themselves have been shown to interact with microtubules via a specific protein, Cellulose Synthase-Interaction 1 (CSI1), an interaction that can also tether CSC-containing organelles and vesicles to microtubules^10–13^. Thus, it was not clear if this exocytotic patterning mechanism may be widely employed in plant cells, or if it is unique to delivery of CSCs to the plasma membrane.

Microtubule targeting of exocytosis is of special interest in the context of plant cell morphogenesis because ordered cortical microtubule arrays are essential to create functional cell shape. Depolymerization of microtubules by oryzalin causes isotropic growth and a complete loss of genetically determined cell shap. The predominant hypothesis for the function of cortical microtubules in directing cell growth is that they guide and organize the trajectories of CSCs as they synthesize and extrude cellulose microfibrils into the cell wall, thus creating material and mechanical anisotropy that constrains and directs expansion of the cell wall (which is under tension generated by high turgor pressure)^14^. We confirmed that individual CSC trajectories are in fact guided by microtubules by live cell imaging studies^15^. However, disruption of both microtubule guidance of CSCs and delivery of CSCs to the plasma membrane in *csi1* null mutants causes only a modest loss of cell growth anisotropy^10,11,13^. In addition, although complete loss of growth anisotropy is caused by microtubule depolymerization, this phenotype is not due to a loss of CSC delivery to the plasma membrane^7^ nor a lack of ordering in CSC trajectories^7,16,17^. Further, cell growth rate does not appear to be impaired by oryzalin treatment and loss of cortical microtubules^18^. These observations suggest that cortical microtubules arrays have additional important functions in establishing cell shape, prompting us to ask if cortical microtubules may guide cellular morphogenesis by acting as a general targeting mechanism, beyond CSC’s, to pattern exocytosis during cell growth.

To determine if individual exocytosis events in elongating plant cells are broadly patterned by cortical microtubule targeting, we require a dynamic biological marker that identifies single exocytotic events and is involved in a wide range of exocytotic traffic to the plasma membrane. To date, such a marker has not been identified. Imaging studies of soluble *N*-ethylmaleimide-sensitive factor attachment protein receptors (SNARE proteins)^19–21^ and components of the exocyst complex^22,23^, have not revealed a clear relationship between the presence and dynamics of these proteins with verified individual exocytosis events nor have they shown association of these proteins with microtubules in growing plant cells^24^. However, Exo70, a core component of the exocyst complex, has been localized to cortical microtubules in differentiating xylem cells^25^, a pattern normally not observed in other cell types^25,26^.

To address this gap in tools for visualizing and studying individual exocytotic events, we investigated use of the essential Sec/Munc (SM) protein KEULE. SM proteins are essential regulators of SNARE-mediated vesicle fusion. In exocytosis, SNARE proteins anchored to the vesicle interact with those on the plasma membrane and form a membrane bridging complex with other partners that enables membrane fusion^24,27,28^. The soluble SM proteins are known to bind to SNAREs, and evidence from genetic studies together with *in vivo* and *in vitro* experiments suggest they function in vesicle docking, SNARE complex assembly and vesicle fusion^29–34^. In plants, the *Arabidopsis* Sec1 homolog KEULE was identified in a genetic screen for embryonic pattern mutants^35^, where loss of function mutations are lethal, causing bloated and multinucleate cells^36^. Consistent with other SM proteins, KEULE interacts genetically and biochemically with Qa-SNAREs, specifically KNOLLE (SYP111) and SYP121, and has been shown to play essential roles in cell plate assembly during cytokinesis and exocytotic traffic to the plasma membrane^19,36–40^. The demonstrated function of KEULE in exocytosis, combined with its severe loss of function phenotype due to lack of exocytosis suggested that KEULE might serve as an excellent candidate for a reporter of bulk exocytotic traffic.

The molecular and structural mechanisms by which SM proteins interact with and regulate SNARE complex assembly have been investigated intensively by genetic, biochemical*, in vitro* and structural studies^29–34^, and SM proteins have been imaged in live cells in several systems^41–48^. However, important features of their behaviors and biological function in living cells remain to be uncovered. These questions include the fine scale cellular distributions of SM proteins and clusters, the association of SM clusters with exocytosis events, the relationship of these exocytosis events to potential organizing machineries and the dynamics of SM assembly and loss at exocytosis sites, especially prior to and during vesicle fusion.

Here, we develop and apply automated and high precision analysis methods to detect and track thousands of individual and fluorescently tagged KEULE clusters to investigate their dynamics and to map their positions at the plasma membrane in living cells. We evaluate these clusters as exocytotic events by use of CSCs as a validated cargo and employ genetic analysis to test for the function of KEULE in CSC delivery. To determine not only whether but what the spatial rules are that govern how microtubules pattern exocytosis, we needed an approach that could reveal probability distributions invisible to the eye in live cell movies. Classical colocalization analysis yields only binary classifications at arbitrary cutoffs, obscuring probability landscapes. We therefore developed a Bayesian inference approach that, combined with maximum likelihood localization of >100,000 events, extracts the comprehensive probability distribution of exocytosis as a function of distance from microtubules. This revealed a non-monotonic spatial profile: flat ∼180 nm enrichment zones flanking microtubules, followed by ∼520 nm depletion zones, with cortical microtubule arrays spatially redistributing ∼13% of KEULE- mediated exocytosis. These zones of enrichment and depletion effectively act as a redistribution and concentrating mechanism to pattern exocytosis in the shape of the cortical microtubule array. Further, the observed probability distribution suggests that a membrane domain model for exocytotic targeting may be more likely than strict microtubule tethering, as the width and flat profile of the enrichment zone would be difficult to reconcile with direct vesicle capture at microtubule surfaces. These results demonstrate that non-centrosomal microtubule arrays in growing plant cells pattern exocytosis through proximity-dependent mechanisms, representing a fundamentally different organizational mode than polarized transport by centrosomal and other polarized arrays, where organization and positioning of plus ends determines targeting.

## Results

### Functional dynamics of tdTomato-KEULE cluster formation

To visualize KEULE protein *in vivo*, we created a tdTomato-KEULE fluorescent reporter constructed from the chromosomal gene with native upstream regulatory sequence and introduced it into *Arabidopsis* plants heterozygous for a *keule* deletion allele. To obtain plants for study of KEULE localization and dynamics, we screened for self-fertilized progeny of these transgenic plants that exhibited rescue of *keule* loss of function phenotype by the tagged KEULE protein. These plants exhibited full rescue of plant growth and development through the 5-day seedling stage used for our further studies (Fig. S1, Methods).

In dark-grown *Arabidopsis* hypocotyl cells expressing the tdTomato-KEULE construct, we observed KEULE signal appear as densely distributed and temporally dynamic puncta at the plasma membrane (Fig. 1A). The dynamics of signal increase and decay were apparent in time-lapse movies (Movie S1), kymographs of signal intensity over time along transect lines (Fig. 1B) and in sequential image samples of individual puncta (Fig. 1C). In all these events, a steady increase in signal intensity was observed, followed by an abrupt decrease and disappearance of signal. This behavior indicates the accumulation and loss of many fluorescently labeled KEULE proteins, which we will refer to hereafter as clusters.

**Fig. 1.**
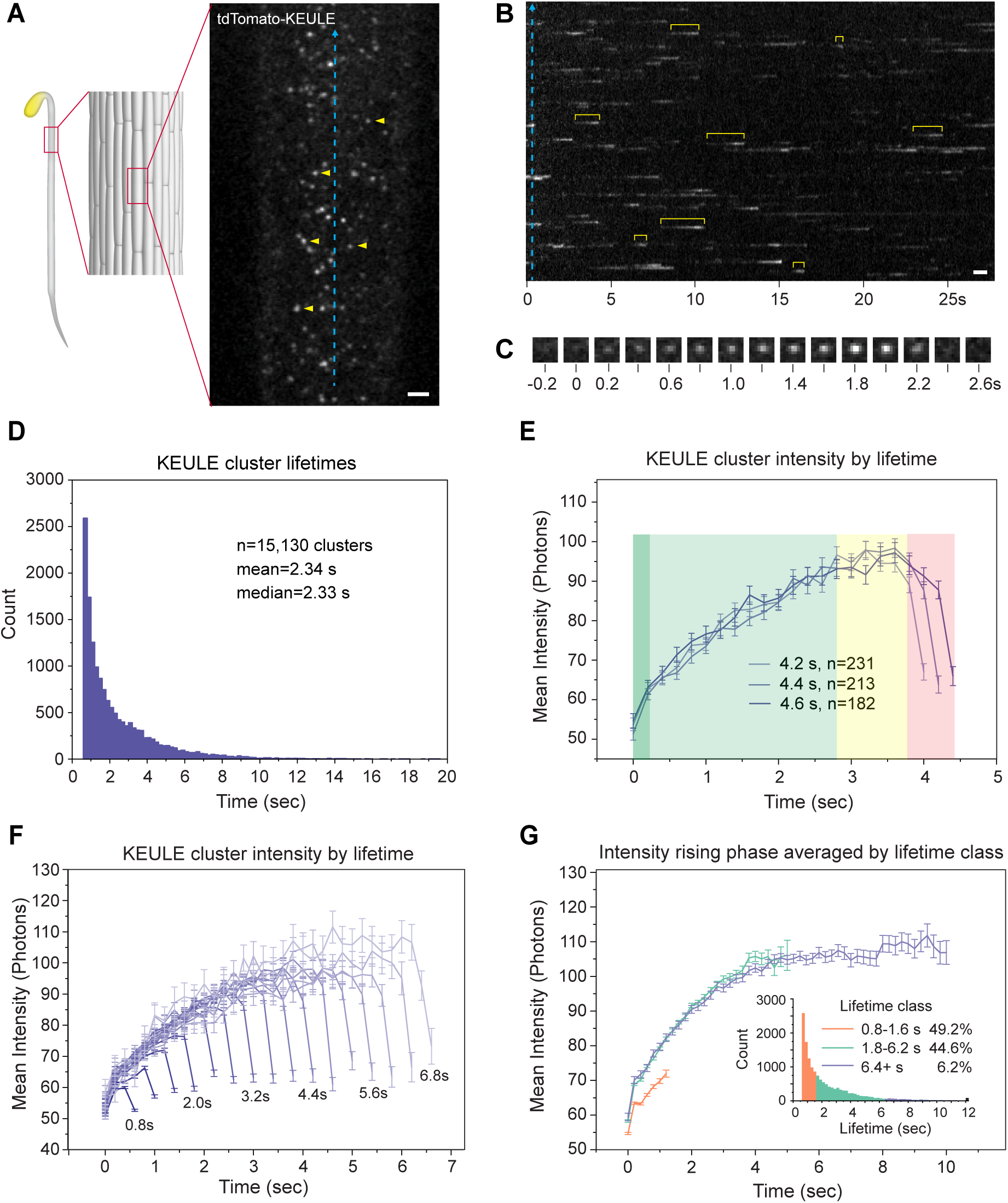
KEULE assembly dynamics at the plasma membrane. (**A**) Distribution of tdTomato-KEULE signal at the plasma membrane of etiolated hypocotyl cells in *Arabidopsis*. Arrowheads point out individual clusters of labeled KEULE protein. (**B**) Kymograph of the tdTomato-KEULE signal intensity over time as sampled over the transect line shown in (**A**) (blue dashes). The yellow brackets highlight several individual tdTomato-KEULE clusters. (**C**) Example of tdTomato-KEULE cluster intensity over time. (**D**) Histogram of KEULE cluster lifetimes in 200 ms bins. n = 15,130 clusters. (**E**) Average tdTomato-KEULE cluster signal intensity with a lifetime of 4.2, 4.4 and 4.6 s. Note the initial rapid increase in signal over the first acquisition step (darker green), the more gradual, near linear, ramp up (green) to a plateau (yellow), followed by abrupt loss to non-detection (red). n = 231, 213,182 clusters for 4.2, 4.4 4.8 s, respectively. (**F**) Average tdTomato-KEULE cluster signal intensity with lifetimes ranging from 0.8 to 6.8 s. n = 15,397 clusters in total, ranging from 4992 clusters for 0.8 seconds to 122 for 6.8 s. (**G**) Average signal intensities of tdTomato-KEULE clusters in three cumulative lifetime ranges, 0.8 to 1.6 s, 1.8 to 6.2 seconds and 6.4+ s. The inset shows the lifetime distribution histogram of tdTomato- KEULE cluster lifetimes with the lifetime ranges color coded. n = 15,130 clusters. All data in panels D-G acquired from 12 cells from 12 seedlings. Error bars are S.E. Scale bars are 2 µm.

The dynamics of KEULE accumulation and disappearance are presumably related to its function. Therefore, we reasoned that a quantitative and robust picture of KEULE cluster dynamics would be useful in gaining insight into the functionality of KEULE clusters and the underlying molecular mechanisms of exocytosis. To accomplish this, we collected a large-scale and high-resolution dataset comprising thousands of individual events. We acquired streamed images at 200 ms intervals and performed automated detection and frame-to-frame linking of KEULE cluster signal using µTrack^49^. These track data were then used to extract position and intensity information by Maximum Likelihood Estimate 2D Gaussian fitting^50^ (see Methods, Movie S2). To define events of signal appearance, accumulation and loss, we used a minimum sample of 4 sequential acquisitions.

The lifetime distribution of measured tdTomato-KEULE clusters showed a large range, from the minimal event length of 800 ms to over half a minute. These data followed a roughly exponential distribution with a mean of 2.34 seconds and ∼95% of lifetimes being less than 7 seconds long (Fig. 1D). The large lifetime range of observed KEULE cluster events might represent a collection of functional classes. If so, such classes may by differentiated by their kinetics. To assess the kinetics of KEULE signal accumulation and loss, and to ask if these kinetics might vary as a function of cluster lifetime, we leveraged our large data set to calculate KEULE kinetics for each 200 ms lifetime increment. Fig. 1E shows average intensities for all observed clusters with lifetimes of 4.2, 4.4 and 4.6 seconds. All these plots show a similar pattern: an initial steep rise in intensity, followed by a slower and near linear increase, a plateau phase that is especially evident in longer events, and ending with a sharp drop in signal. If the clusters represent exocytotic events, these phases may reflect the recruitment of KEULE and assembly of the exocytotic apparatus, a maturation phase, and finally rapid loss of KEULE as the cytokinetic apparatus is disassembled upon membrane fusion.

To quantitatively compare differences in accumulation kinetics as a function of cluster lifetime, we grouped events into three duration classes; short (0.8 to 1.6s), medium (1.8 to 6.2s) and long (6.4+s) – and then averaged the intensities at each 200 msec timepoint (Fig. 1G, for details see Methods). The timepoints following the plateau phase were omitted to aid comparison of signal accumulation trajectories. This analysis revealed that, KEULE signal, and thus KEULE protein, accumulates at slower rate in short-lived events as compared to medium- and long-lived events, which displayed nearly identical trajectories of signal accumulation (Fig. 1G). These data suggest that short and longer lived KEULE clusters may differ functionally in regard to KEULE recruitment. It is noteworthy that for events up to ∼6 seconds, the maximum signal intensity achieved increased steadily as a function of cluster lifetime. SEM studies of *in vitro* vesicle fusion have shown that more SNARE proteins are present at the fusion sites of larger vesicles as compared to smaller vesicles^51,52^. As SEC/MUNC proteins interact with SNARES, it is therefore possible that events with longer and high levels of KEULE represent exocytosis events of larger vesicles. In events lasting longer than about 6 seconds, maximum signal no longer increased with lifetime but instead showed a prolonged plateau phase, possibly indicating stalled exocytosis events or alternatively, an extended checkpoint for exocytosis.

### Clusters of tdTomato-KEULE coincide with CSC delivery to the plasma membrane

To determine if dynamic KEULE clusters are associated with exocytosis events, we asked if they were associated with the delivery of exocytosis cargo. To perform this experiment, we took advantage of our previous work on delivery of CSCs to the plasma membrane^7^. CSCs are assembled in the Golgi system from multiple CESA catalytic subunits^53,54^, move through the trans-Golgi network (TGN) and are budded off into vesicles that fuse with the plasma membrane^7^. Because CSCs are formed of multiple CESA subunits, they can be labeled by fluorescent proteins in sufficient number to allow their detection and the tracking of individual complexes during the process of exocytosis. Labeled vesicles display rapid and erratic dynamics that abruptly become stabilized at the optical plane of the plasma membrane with a mean duration of about 60 seconds, followed by a transition to slow, steady and typically linear motility of between 250 and 450 nm/min. The transition erratic to static movement is interpreted as evidence for vesicular capture and tethering at the plasma membrane, and the transition to slow and steady movement indicates successful vesicle fusion and delivery of CSCs, which are propelled by the synthesis and extrusion of cellulose microfibrils^7,15^. This stereotyped behavior makes CSCs an excellent reporter for successful exocytosis, revealing a time and place for vesicle tethering, and distinguishing vesicle tethering at the plasma membrane from successful vesicle fusion and cargo delivery.

If KEULE is involved in the exocytotic delivery of CSCs to the plasma membrane, we expect to observe co-localization between CSC delivery events and KEULE clusters. To visualize KEULE together with CSCs, we generated plants that express tdTomato-KEULE and CIT-CESA6, a catalytic CSC subunit labeled with Citrine fluorescent protein^15^ in a *keule*/*cesa6* double mutant background (Methods). As CIT-CESA6 signal is relatively dense in the plasma membrane of rapidly growing cells, we photobleached the CIT-CESA6 signal over a region of the plasma membrane to aid detection of new CSC arrivals and their relation to td-Tomato-KEULE signal (Fig. 2A, Movie S3). For each CSC arrival at the optical plane of the plasma membrane, we determined the first frame in which the CESA6 signal became stationary (an indication of interaction and tethering at the plasma membrane). We then identified the nearest KEULE cluster in a range of −10 to 50 seconds of the first stable CESA6 frame and measured the distance between the centroids of the CESA6 spot and the nearest KEULE spot by estimating their coordinates using maximum likelihood 2D Gaussian fitting (See Methods, Fig. 2B and D). A search range starting at −10 seconds to the estimated tethering time was chosen because the exact frame of stationary transitions was in some cases obscured by a variable background of other labeled organelles such as Golgi bodies. To assess statistically the spatial and temporal association of CSC delivery events and KEULE clusters, we generated a simulated dataset of random locations for CSC delivery within the photobleaching areas and identified the nearest KEULE spots to these random locations within the same time window we used for our experimental observations. We then compared the distributions of distances to the nearest KEULE cluster for the observed and simulated datasets and found that the experimental data were strongly skewed to shorter distances ((Fig. 2B, p < 1E-06, Kolmogorov-Smirnov test). The pattern of over- and under- representation of observed with respect to simulated data is more readily seen in a plot of the bin-by-bin differences, where shorter KEULE-microtubule distances are seen to be over-represented in bins below 200 nm and underrepresented in bins of longer values (Fig. 2C). Using a Bayesian approach (see Methods), we found that the most likely values of the experimental biases from the null hypothesis of a random association were significantly above that expected by chance alone for distance bins below 120 nm (Fig 2E and F) and significantly lower than expected in bins from 200 to 480 nm. Based on these data we defined colocalization as observing a KEULE cluster centroid less than 200 nm from the position of CSC signal in the first frame in which it was observed to stabilize (and in a time window of −10 to 50 seconds around that frame). By these criteria, colocalization occurred in 81% (157 out of 194) of observed delivery events compared to 29% (13035 out of 45051) of random locations at the plasma membrane (p < 1.4E-50, Fisher’s exact test). Thus, these data indicate that KEULE clusters were observed to be significantly associated with the great majority of CSC deliveries.

**Fig. 2.**
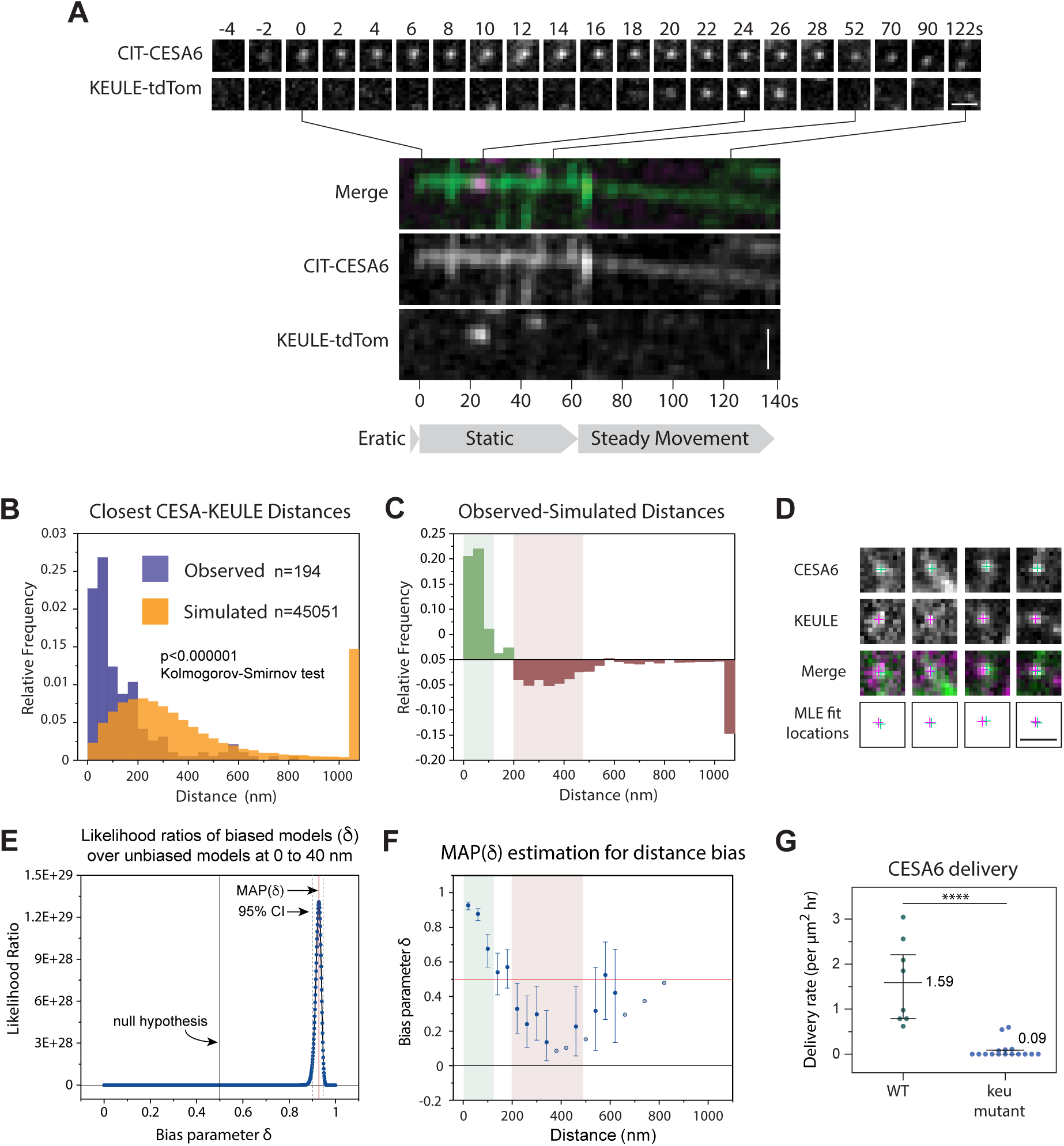
KEULE is spatially associated with and required for CSC delivery to the plasma membrane. (**A**) CESA exocytosis event. (Top panel) Time series (top panel) and kymograph (bottom panel) of tdTomato-KEULE and CIT-CESA6 during a CESA exocytosis event. The kymograph shows evidence for CSC-vesicle tethering at the plasma membrane (start of static phase) and successful CSC delivery to the plasma membrane as an active complex (steady movement phase, see text). (**B**) Distribution of distances between the positions of CIT-CESA6 signals at start of static phase and the nearest tdTomato-KEULE cluster, as estimated by MLE PSF fitting. Data shown both for observed CIT-CESA6 locations and simulated random locations. (**C**) Differences of the observed vs expected CESA6-KEULE distance frequencies in (**B**), portraying over- and under-representation (green and red respectively). (**D**) Examples of colocalization and fitted positions for events in (**B**). (**E**) Bayesian likelihood ratios of the observed frequency of distances between 0 and 40 nm given bias δ over δ(0.5). The position of the peak value indicates the most likely, or maximum a posteriori probability (MAP), value for the bias δ for the observed frequency given the expected frequency. (**F**) Bayesian analysis of the MAP(δ) for each distance bin, MAP(δ) = 0.5 indicates no difference from expected (indicated by red line). Error bars are 95% confidence intervals expressed as the highest posterior density (HPD, the shortest interval containing 95% of the probability density). (**G**) CESA6 delivery rates in wild type and *keule* mutant cells. Whiskers are quartiles. **** p<0.0001 Wilcoxon 2-sample test. In panels B-F, n = 194 CESA exocytosis events in 8 cells from 8 seedlings for observed data. For simulated data, n = 45051 randomized CESA exocytosis locations in the same KEULE channel data sets. In panel G, n = 8 cells from 8 seedlings for WT and 16 cells from 16 plants for *keule*. Scale bars are 1 µm.

We considered that our automated experimental protocol may underestimate colocalization between CSC delivery and KEULE. For example, the time interval used for these experiments was 2.11 seconds, of which 400 ms was used for tdTomato-KEULE excitation and image acquisition. Since a substantial portion of KEULE cluster lifetimes is shorter than the ∼1.7 second gaps between image acquisitions, we do not expect to detect all KEULE clusters within the −10 to 50 second experimental window. In addition, not all excited KEULE foci are successfully identified by the spot detection and filtering routines used for large-scale data collection. To understand how these two cases, no excitation and no detection, affect our colocalization results, we performed a hand-curated analysis using a custom MATLAB GUI. For each CSC delivery, we examined whether there was overlap of KEULE signal with the centroid of the CSC signal in the first frame of positionally stable KEULE signal (tethering frame). We found that in 92% (159 out of 172) of the cases, KEULE signal was observed to overlap the CSC position in the tethering frame within −10 to 50 seconds of CSC stabilization, suggesting that our automated analysis, while free of possible observer bias, likely underestimates the true degree of colocalization frequency of CSC exocytosis events and KEULE.

If KEULE clusters function in CSC exocytosis after vesicle tethering, we thought it likely that not only should they colocalize with CSC signal, but that the time intervals between vesicle tethering and the appearance of KEULE should be constrained by mechanisms of specific KEULE recruitment and have a non-random distribution. We tested this prediction by comparing the delay times for arrival of the nearest KEULE spot following the tethering frame in the observed vs the simulated KEULE data sets. The observed delay time distribution was significantly different from the random distribution of the simulated data (p < 7.8E-04 Mann-Whitney U test) (Fig. 3A), having a peaked distribution with a median of 2.111 seconds and a mean of 7.92 s, suggesting that KEULE is recruited and functions within several seconds after the CSC vesicle is stabilized and tethered at the plasma membrane.

**Fig. 3.**
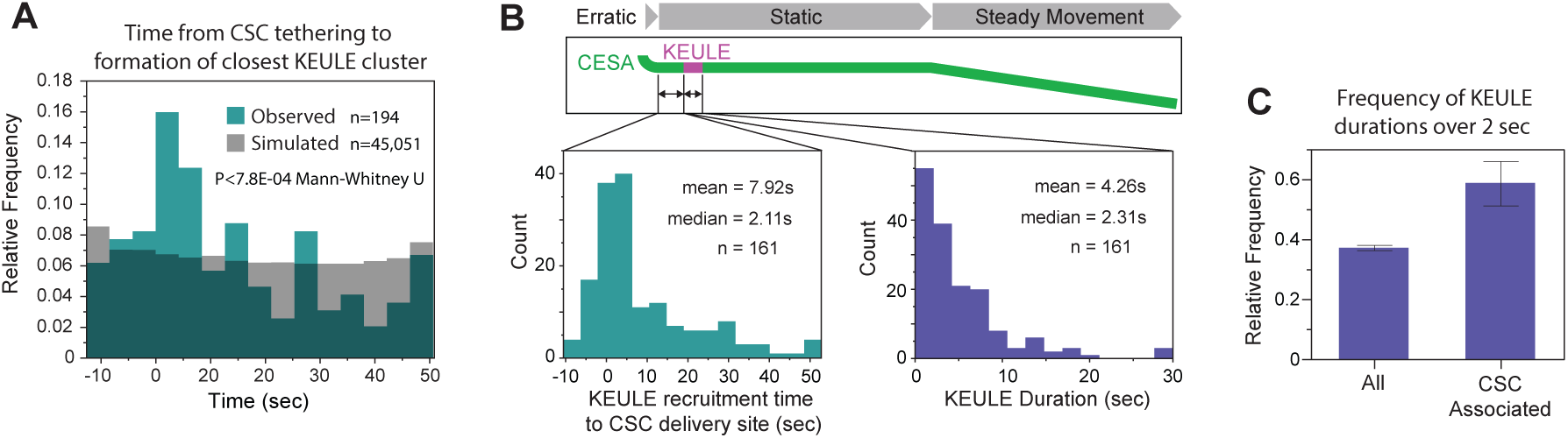
KEULE cluster appearance is temporally associated with CSC appearance at the plasma membrane. (**A**) Comparison of the waiting time distribution from CSC signal stabilization until the detection of the nearest tdTomato-KEULE cluster in observed and simulated data. n = 194 CSC delivery events in 8 cells from 8 plants. (**B**) KEULE dynamics in the context of CSC delivery to the plasma membrane. Histograms of waiting times (left pane) and lifetimes of tdTomato-KEULE clusters co- localized with CSC delivery events (right pane). n = 161 CSC delivery events in 8 cells from 8 plants for observed data, n = 45,051 simulated CSC delivery events applied to the same data sets. (**C**) KEULE cluster durations for all events and those associated with CSC delivery, compared by relative frequency of durations over 2 seconds. n = 13766 for all KEULE clusters, n = 180 for KEULE clusters associated with CSC delivery. Whiskers are exact binomial 95% confidence intervals, p = 6.1E-09 Fishers’ exact test.

Our measurements of KEULE cluster dynamics revealed a wide range of lifetimes with similar kinetics of signal gain and loss. Are exocytotic events associated with both long and short KEULE cluster lifetimes? The distribution of KEULE cluster lifetimes associated with CESA deliveries was similar to that for the general population, with both having a similar range of values and being roughly exponential with a long tail (Fig. 3B compared to Fig.1D). However, while similar, the distribution of lifetimes associated with CESA delivery was observed to have a bias towards longer values (median 4.3 seconds, as compared to 2.3 seconds for clusters in general). To test the significant of this bias, we compared the relative frequencies of durations longer than two seconds and found a highly significant difference - 59% vs 37% for CESA delivery-associated KEULE clusters events vs clusters in general (Fig. 3C, p = 6.07E-09 Fisher’s exact test). These results indicate that both short-lived and long-lived KEULE clusters are associated with verified exocytosis events and that those associated with CESA complexes are modestly biased towards longer lifetimes. Taken together, our data provide insight into the kinetics of SM protein recruitment to sites of exocytosis and reveal the time of vesicle fusion following the stabilization and tethering of exocytotic vesicles at the plant cell plasma membrane.

Our data show that KEULE clusters are associated with confirmed sites of CSC exocytosis in both space and time, but is KEULE function required for CSC exocytosis? To answer this question, we quantified CSC delivery in selfed progeny of parental plants homozygous for a transgene expressing CIT-CESA6, heterozygous for a *tdTomato-KEULE* allele, and homozygous for *cesa6* and *keule* loss of function alleles (See Methods). Seedlings tested for loss of KEULE function were identified by morphological *keule* phenotype (severely reduced in size, large and misshaped cells) and little to no tdTomato-KEULE signal. In these mutant seedlings, we found that the CSC delivery rate was reduced by over 17-fold as compared to that in wildtype (p<0.0001, Wilcoxon two sample test, Fig. 2G). Taken together our data show that KEULE clusters are spatially and temporally associated with, and functionally required for, the successful exocytosis and delivery of CSCs to the plasma membrane. These results indicate that KEULE clusters of stereotyped dynamics are functional markers of exocytosis events.

### KEULE dynamics are sensitive to drugs and treatments that reduce exocytosis

We previously reported that the number of active CSCs in the plasma membrane were rapidly and significantly reduced by treatment with mannitol (osmotic stress) and the herbicide isoxaben^7^. Measurements of CSC delivery revealed that mannitol treatment also caused a 100-fold reduction in the delivery rate of CSC’s, indicating that CSC loss at the plasma membrane was likely caused primarily by inhibition of exocytosis. Likewise, isoxaben has been characterized as a specific cellulose synthase complex inhibitor^55^, but its mode of action is also not well understood. Here we (1) determine if isoxaben treatment also reduces CSC exocytosis and (2) ask if KEULE dynamics may provide insight into how osmotic stress and isoxaben act to inhibit exocytosis.

To address the above questions, we treated seedlings expressing tdTomato-KEULE and Cit-CESA6 with either 200 mM mannitol, 100nM isoxaben or carrier only (0.01% DMSO, mock) and assayed CSC delivery to the plasma membrane as described above. We found that, like osmotic stress caused by mannitol, isoxaben is also a potent inhibitor of CSC exocytosis, reducing CSC delivery rate by ∼70 fold (Fig. S2). By contrast to CSC delivery, the overall frequencies of KEULE cluster formation after both mannitol and isoxaben treatment were not significantly lower than that observed for mock treatment, indicating that neither acts to absolutely block initial KEULE recruitment and cluster formation rates (Fig. S3A). However, when we examined KEULE accumulation dynamics we observed striking differences among treatments (Fig. S3B, C). In cells treated with mannitol KEULE signal accumulated more slowly and to lower levels in the short and intermediate lifetime classes events (Fig. S3B). Intriguingly, the trajectories of long duration clusters (6.4 seconds or longer) exhibited a markedly different pattern. KEULE signal trajectories for both mock and mannitol treated cells tracked each other closely for the first 5 seconds or so, but rather than reaching a plateau after 5 seconds, signal in mannitol treated cells continued to increase steadily until the transition to rapid signal loss (Fig. S3B). Taken together, these data suggest that there may be two different mechanisms by which mannitol treatment and osmotic stress affects KEULE dynamics, depending on their duration class: one mechanism that acts negatively and from cluster inception that reduces the rate and level of KEULE accumulation in short and intermediate events, and one which acts at a later stage in long duration events that prevents transition to the normally occurring plateau stage. By contrast, we observed that isoxaben treatment slowed accumulation of tdTomato-KEULE across events of all durations (Fig. S3C).

We observed related differences among treatments in KEULE cluster lifetimes. In comparing the distributions of cluster lifetimes in mock and mannitol treated cells, a bimodal effect was again observed, with both shorter and longer lifetimes being over-represented in comparison with mock treatment (and intermediate events being under-represented) (Fig S4A, B, D). As was the case with intensity dynamics, isoxaben treatment showed more of a unimodal effect, with the lifetime distribution being shifted to the left (Fig. S4B, C, E). Mannitol and isoxaben also caused diverging effects on persistence of KEULE clusters, as assessed by survival analysis (Fig. S4F, G). Taken together, our data show that KEULE cluster dynamics are sensitive to perturbations that affect exocytosis and reveal evidence that mannitol and isoxaben treatments affect these dynamics in distinct ways, suggesting they may perturb exocytosis by different mechanisms.

### Cortical microtubules pattern KEULE cluster distribution and exocytosis

The measured flux of KEULE cluster events was 85-fold higher than that measured for CSC delivery (144 events per µm^2^ per h, 1.59 evens per µm^2^ per h). Having established dynamic KEULE clusters as markers of exocytotic events, we now have a tool to test if a major class of exocytotic traffic was non-randomly distributed at the plasma membrane with respect to the cortical microtubule cytoskeleton. Further, the ability of our methods to map the probability of exocytosis as a function of the distance to microtubules allowed us to ask questions about the biophysical mechanisms of targeting. For example, does the frequency of microtubule directed exocytosis peak at the estimated positions of microtubules, such as would be expected if exocytotic organelles interact directly with them, or might

To answer these questions, we visualized KEULE cluster events together with cortical microtubules, by performing live cell imaging in hypocotyl cells of *Arabidopsis* seedlings expressing GFP-TUA5 and tdTomato-KEULE in a *keule* knock-out background (Fig. 4A, Movie S4). Acquiring images at one second intervals, we tracked individual KEULE events at the plasma membrane and used MLEwG fitting to extract information about their precise positions (Fig. 4B, Methods). To determine the location of the nearest microtubule to each detected KEULE cluster, we identified the frame with peak signal intensity as the reference point for cluster position (Fig. 4C panels 1-3) and measured signal intensity in the microtubule channel along a series of scan lines that were rotated at 9° intervals around the centroid of the KEULE cluster (Methods and Fig. 4C panel 4). On each of these scan lines, we detected all microtubule signal peaks and fit them with a modified quasi-1-dimensional MLEwG fit (Methods, Fig. 4D). Since the microtubule signal is best represented as a quasi-1-dimenional Gaussian orthogonal to the microtubule axis, we determined the orientation of the nearest microtubule as identified by peak analysis and used the scan line most orthogonal to the microtubule angle for determining the location of the microtubule central axis closest to the KEULE spot in question (Fig. 4C, panel 5). For example, the intensity of KEULE signal in photoelectrons for the most orthogonal scan line in Fig. 4C panel 5 is shown in Fig. 4D panel 1, the triple Gaussian 1D MLEwG fit in panel 2, and the decomposed individual peak fits in panel 3. We used the fit positional coordinates of the nearest microtubule signal peak and the KEULE cluster to calculate the distance and the photoelectron fit values to estimate the standard error of the measurement (See Methods and Fig. 4C panel 6). We repeated this process for all detected KEULE tracks in each movie to assess the spatial relationship between KEULE clusters and the nearest microtubules to those clusters (Fig. 4E, panel 1).

**Fig. 4.**
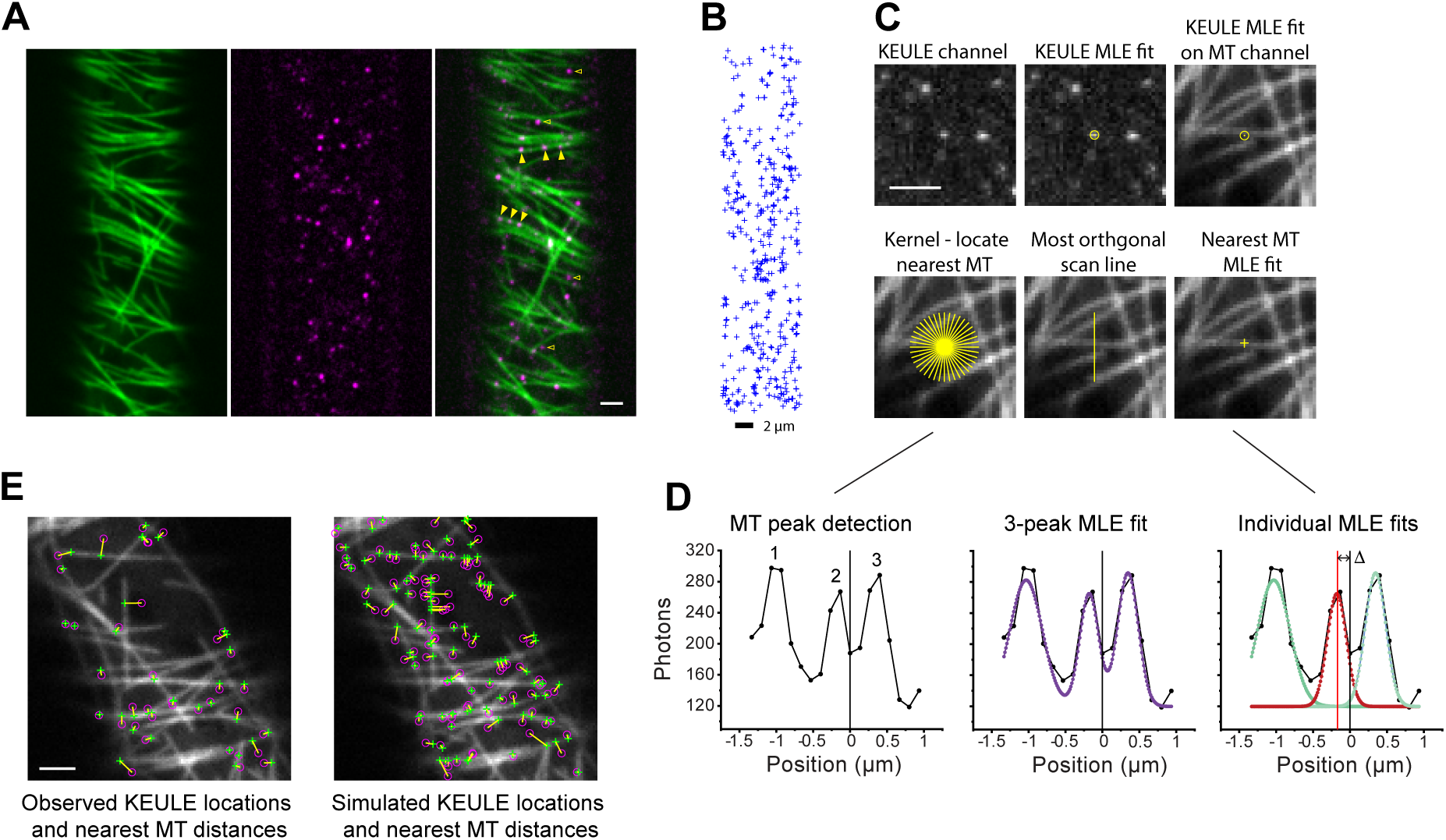
Microtubule dependent distribution of exocytosis events. (**A**) Images of GFP-TUA5 and tdTomato-KEULE signal in epidermal cells. Arrowheads indicate examples of apparent co-localization of signal. (**B**) Spatial patterning of tdTomato-KEULE clusters over the range of 36.8 seconds. Crosses show positions of MLEwG fits. (**C**) Steps in the routine employed to calculation the distance from a KEULE cluster to the center of the nearest microtubule distance. (**D**) Steps in quasi-1-dimensional MLEwG fitting of microtubule signal along the most-orthogonal scan line of a rotating filter. (**E**) Overlay of a microtubule channel image and pairs of markers for MLE fit locations of KEULE clusters and the center of the nearest microtubule in observed data (left) and for simulated random KEULE locations (right). KEULE cluster detects from −5 to + 5 frames from the microtubule image are shown. Scale bars are 2 µm.

To ask if the measured distribution of KEULE to microtubule distances differs from that expected if KEULE cluster positions are independent of microtubule locations (the null hypothesis), we first needed to determine what the expected distributions are under the null hypothesis when our detection and analysis routines are applied to the image datasets we used for our experimental observations. This is necessary because microtubule array structure, which varies within and among cells, will have a strong influence on the distributions of distances, as will the dimensions of our detection kernels. Therefore, we generated a simulated set of random locations within each region of interest across all image frames in the movies and calculated the distances from these synthetic KEULE foci to the nearest microtubule (Fig. 4E, panel 2). When the experimental and expected distributions were plotted together as relative frequencies, they showed evidence for distinct differences (Fig. 5A). The spatial pattern of under- and over- representation is more easily visualized in a plot of the differences between observed and expected frequencies (Fig. 5B). To quantify over- or under-representation of the experimental data given the expected data under the null hypotheses, we used a Bayesian method (See Methods) to determine the most likely positive or negative bias at each distance bin (Fig. 5C and D and Methods). We found that at KEULE-microtubule distances smaller than 180 nm, the maximum likelihood bias was significantly higher than expected from random, while between 220 and ∼700 nm this bias was significantly lower than expected from random. At distances > 700 nm we again found a positive, but smaller, bias (Fig. 5D).

**Fig. 5.**
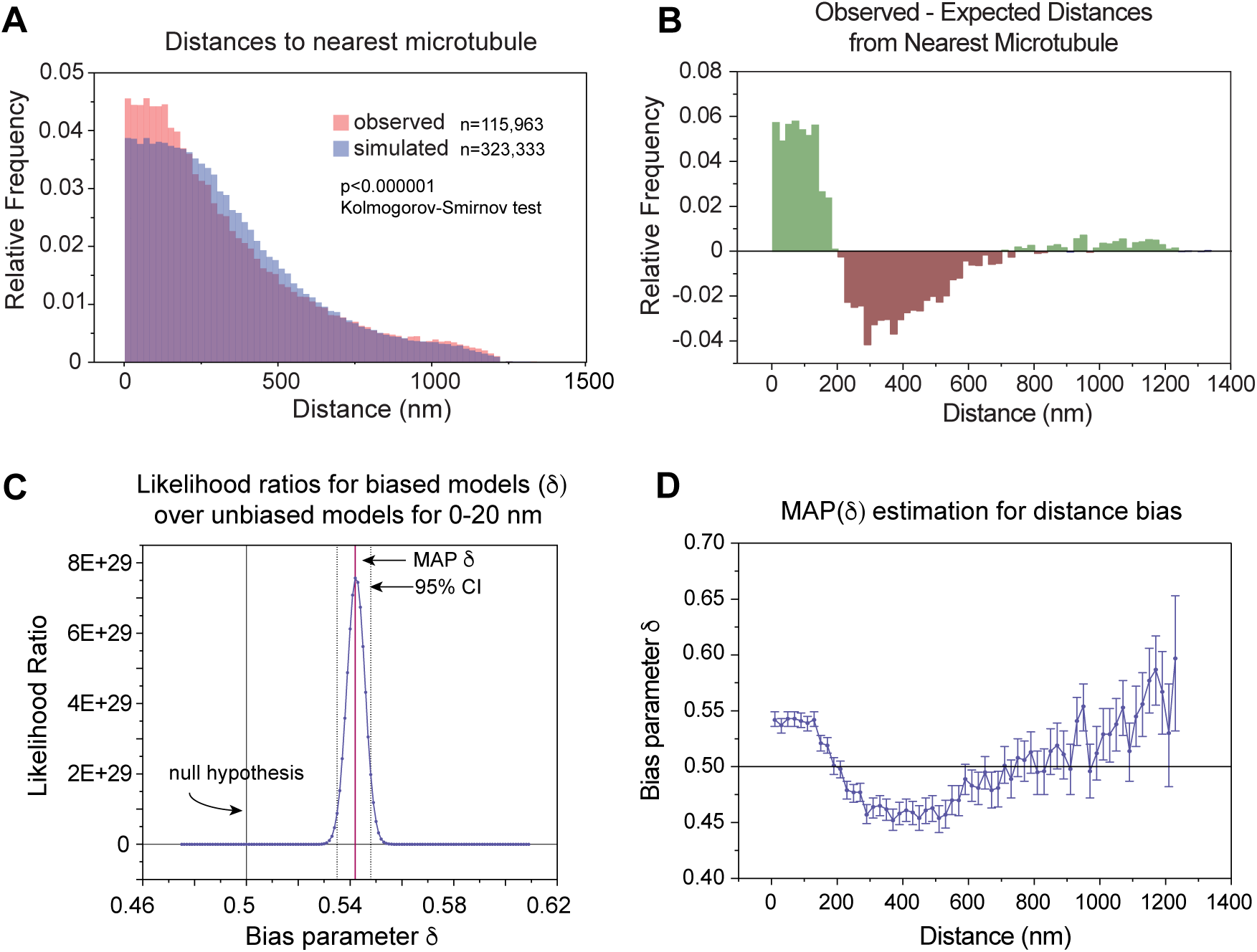
Bayesian analysis of microtubule bias for exocytosis events. (**A**) Comparison of the distributions of KEULE-nearest microtubule distances for observed and simulated data. n = 115,963 KEULE clusters in 14 cells from 14 plants. (**B**) Differences of the observed vs expected KEULE- microtubule distance frequencies shown in (**A**), portraying the pattern of over- and under-representation (green and red respectively). (**C**) Bayesian likelihood ratios of the observed frequency of KEULE- microtubule distances between 0 and 20 nm given bias δ over δ(0.5). (**D**) Bayesian analysis of the MAP(δ) of each distance bin. MAP(δ) = 0.5 indicates no difference from expected (indicated by black line). Data acquired from 14 cells in 14 seedlings. n = 115,963 KEULE clusters in 14 cells from 14 plants. Error bars are 95% confidence intervals expressed as HPD.

Taken together, our results provided strong evidence that exocytosis mediated by KEULE, the major SEC/MUNC protein in *Arabidopsis*, is not spatially random but is organized by microtubules through increasing the frequency of KEULE-mediated exocytosis within a distance of ∼180 nm from their centers. The width of the adjacent zone of negative maximum likelihood bias was observed to be approximately three times wider. Therefore, the measured spatial distribution of enrichment and depletion bias effectively acts as a concentrating mechanism for KEULE-mediated exocytosis that is patterned by cortical microtubule organization.

Two addition features of the spatial patterns of exocytosis bias stood out in particular, raising questions about the underlying classes of biophysical and molecular mechanisms. First, the width of the proximate zone of increased bias was quite large with respect to the scale of microtubules (∼25nm for a single polymer and ∼ 75 for a bundle of two) and the range of action that might be expected for mechanisms involving simple protein linkers between microtubules and “typical” exocytosis vesicles (∼70-90 nm). Second, the estimated values of positive maximum likelihood bias were flat from the closest distance bin of 20 nm all the way to 180 nm, before dropping sharply to negative values. In a mechanism that is sharply focused by microtubules, you might expect bias to peak at the microtubule location and fall off monotonically with distance. We present two possible models to account for these observations, as depicted graphically in Figure 6 and discussed below.

**Fig. 6.**
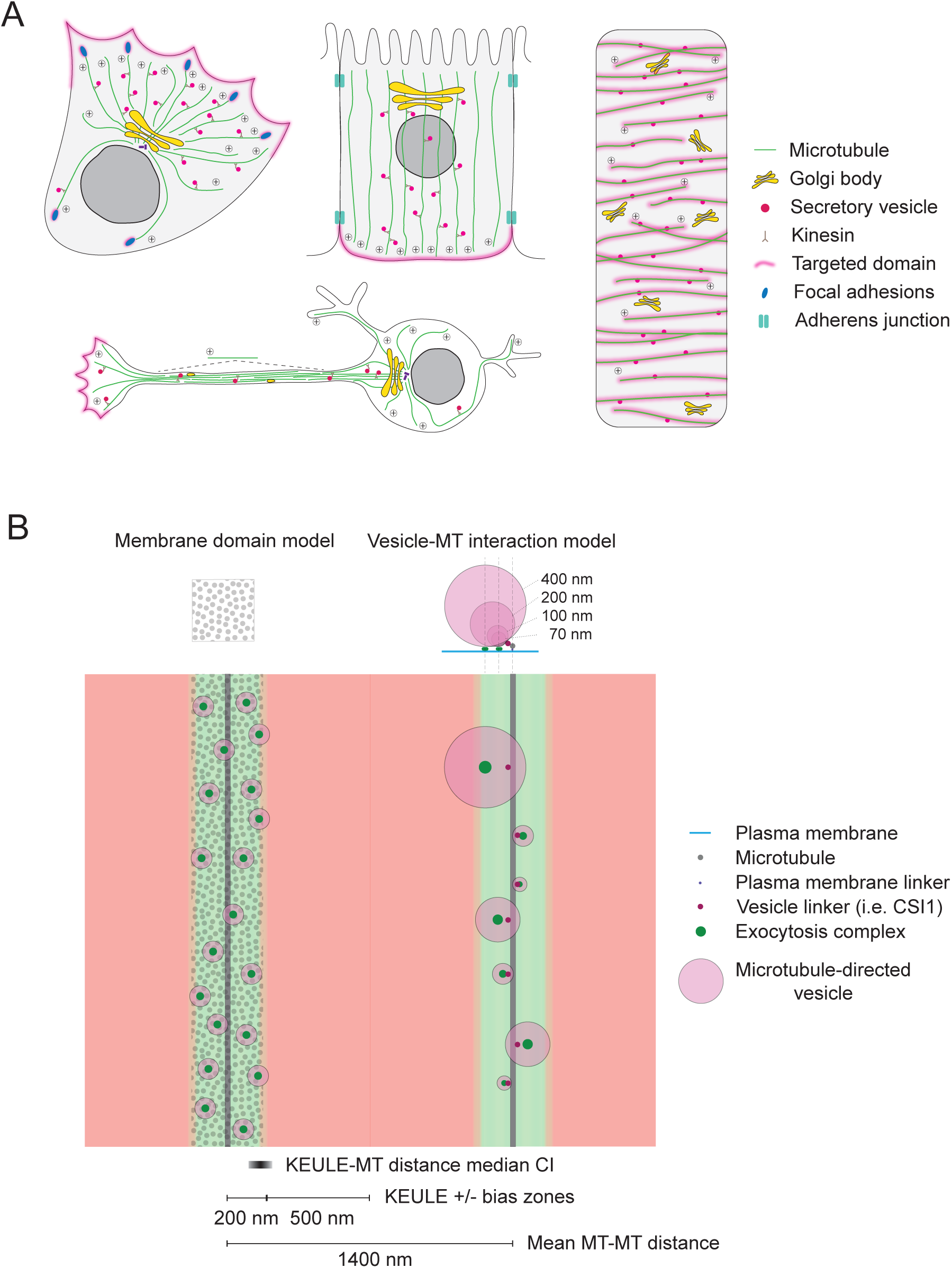
Patterning of exocytosis by microtubule arrays. **(A)** Schematic diagrams of microtubule- directed patterning of exocytosis. The three cells on the left show targeting of exocytosis by motor-driven transport of vesicles along polarized microtubule tracks to locations defined by their plus ends^3^ . Clockwise from upper left: migrating fibroblast, epithelial cell, neuronal cell. By contrast, the diagram at far right shows an epidermal cell in the plant axis, where a 2-D array of microtubules at the cell cortex acts as a template that directs exocytosis in a complex but ordered pattern. The organization of the microtubule lattices defines this pattern. **(B)** Alternative models for spatial patterning of exocytosis by cortical microtubule arrays in plant cells. All structures are abstracted but depicted at approximate scale. Microtubules (grey) are shown spaced at the mean spacing that has been measured in the cortical arrays of growing cells in the hypocotyl epidermis of dark grown *Arabidopsis* seedings^61^. The likelihood of exocytosis marked by dynamic KEULE clusters is shown as a red-green heat map as scaled from the observed over-representation and under-representation of KEULE clusters compared to that expected from the null hypothesis of a random distribution (depicted in Fig. 5B). The two sides of the diagram depict alternative models to explain the measured spatial dimensions of the domains enriched in exocytosis. On the right, the measured extent of the enriched domain is determined by the interaction exocytotic vesicle with microtubules, where those vesicles have a wide range of sizes. On the left, microtubules establish a membrane domain that extends approximately 180 nm on either side where proteins needed for vesicle capture or exocytosis are concentrated. These models are not mutually exclusive. The grey-scale bar at lower right depicts the mean 95% confidence interval for the measured distances of KEULE clusters to microtubule centers.

## Discussion

To date, no native probes for visualizing, measuring and mapping individual exocytosis events in plant cells have been reported other than genetically tagged CSCs, which are a minor and likely specialized part of exocytotic traffic. Here we explored KEULE, considered the major SEC/MUNC mediating exocytosis in Arabidopsis^37–39^, as a candidate for such a tool. The established function of SEC/MUNC proteins in SNARE complex formation and membrane fusion, together with our quantitative evidence that KEULE clusters of stereotyped dynamics are tightly associated with and required for CSC exocytosis, indicate that imaging of fluorescently tagged KEULE can be used to map and investigate the cellular mechanisms that organize and pattern exocytosis in plant cells.

Our measurements of thousands of KEULE clusters at the plasma membrane allowed us to analyze patterns of signal accumulation and loss as a function of varying cluster duration - revealing features of, and raising new questions about, the underlying mechanisms of exocytotic machinery assembly, maturation and resolution. KEULE recruitment features distinct phases: initial rapid accumulation, slower near-linear accumulation, a plateau phase, and abrupt signal loss. The linear accumulation phase suggests constant-rate assembly, potentially consistent with sequential addition of SM-bound SNARE complexes to a growing chain^56–58^. The correlation between assembly duration and peak signal intensity may reflect variation in vesicle size, as larger vesicles recruit more SNARE proteins in vitro^51,52^, The plateau phase likely represents a maturation or checkpoint state preceding membrane fusion.

We found that KEULE dynamics were changed when we disrupted exocytosis with osmotic stress by treatment with mannitol, or by application of a drug, isoxaben. The changes in dynamics were qualitatively different for the two antagonists, indicating that they likely act on different processes affecting exocytosis. These observations indicate that KEULE dynamics, which likely indicate functional aspects of vesicle fusion, are a sensitive and useful readout, or marker, for studying exocytosis and the action of proteins, drugs or physiological conditions that disrupt or alter exocytosis.

High resolution mapping and Bayesian analysis of individual exocytosis events marked by KEULE clusters revealed that these events are not randomly distributed in growing hypocotyl cells but are significantly over-represented in domains defined by the cortical microtubule array. This Bayesian inference approach was essential for revealing probability distributions invisible in live cell imaging data, where classical colocalization methods would provide only binary classifications at arbitrary distance cutoffs. By combining maximum likelihood localization with Bayesian estimation of bias parameters across distance bins, we extracted the comprehensive probability distribution of exocytosis as a function of microtubule proximity, revealing a non-monotonic spatial profile that would be obscured by conventional analysis methods. This mode of patterning of exocytosis differs from that previously described by microtubule arrays that are organized by centrosomes or spindle pole bodies. Microtubules in the latter arrays facilitate and organize exocytosis by directionally transporting proteins and vesicles via molecular motors from the cell body to locations in the cell periphery. The patterning of these destinations is therefore shaped by the polarized organization of these arrays and the spatial positioning of microtubule plus ends. By contrast, our data present quantitative evidence of a different mode of exocytotic organization as determined by non-centrosomal cortical arrays. In these arrays, microtubules are planar and are not polarized by an organizing center. Rather than defining oriented paths from sites of cargo production to cargo delivery, the cortical array as a whole acts as a two-dimensional template to pattern a subset of exocytosis as an image of itself (Fig, 6A). Cortical arrays are frequently highly organized, ordered and specifically rearranged by external and internal cues that alter or control cellular growth, thus exocytotic patterning by these arrays can be dynamically steered. As discussed further below, such highly organized and dynamic patterning of exocytotic traffic may be useful for creating functional and mechanical asymmetries to support crucial functions like morphogenesis.

Our measurements of both microtubule and KEULE cluster locations using sub-resolution MLE fitting, combined with Bayesian analysis of the distributions of the measured distributions of distances, provided quantitative information about the spatial extent of microtubule influence on exocytosis, the bias of that influence as compared to a random distribution, and how that bias varies as a function of distance from microtubules. These data yielded a variety of new insights into how microtubules shape exocytosis. First, these data revealed that microtubules influence exocytosis both positively and negatively and do so over spatial domains much larger than the scale of microtubules and microtubule bundles. The combined width of over-represented and under-represented domains flanking the sides of microtubules and microtubule bundles was measured to be ∼700 nm (Fig. 6B).

Second, the combination of narrower elevated domains coupled with wider depleted domains of elevated exocytosis frequency suggests a concentrating effect of approximately three-fold that further enhances the relative difference in the portion of exocytosis traffic over-represented adjacent to microtubules as compared to distances several hundred nanometers away. Vesicles targeted by microtubules might capture local KEULE associated with the plasma membrane, perhaps as bound to SNARE partners such as SYP121^37^, depleting KEULE concentration in the nearby plasma membrane and making exocytosis there less likely, thus creating the observed zones of reduced exocytosis. Our analysis indicates that ∼13% of KEULE associated exocytotic events are redistributed by microtubules. This represents a significant but partial spatial reorganization of exocytotic traffic, where microtubules create order without absolutely determining all exocytotic sites. The combination of ∼180 nm enrichment zones and ∼520 nm depletion zones flanking each microtubule creates a spatial filter that, given the typical ∼1.4 μm spacing of cortical microtubules in these cells^15^, influences exocytosis across most of the plasma membrane (Figure 6B).

Third, the estimated values of positive bias were remarkably flat all the way from the first distance bin distance bin to 180 nm outboard, after which they dropped sharply to a negative bias. The very high likelihood ratios and narrow confidence windows for these bias estimates, together with the abrupt transition to lower values after 180 nm, suggest that the evidence for the flat character of the bias over this region is strong. We were struck by both the size and shape of this positive bias domain. The width of the positive bias domain is large with respect to the scale of relevant molecules and organelles including microtubules, possible linker proteins and the sizes of typical exocytosis vesicles. Further, if microtubules are targeting exocytosis by direct action of microtubules, such as by a linker protein, estimates of bias might be expected have peak at or near the microtubule then fall off monotonically with distance. Instead, bias appeared to be biphasic.

In figure 6B, we present two alternative models that might explain the extent and pattern that we observed for bias for exocytosis associated with cortical microtubules. In the first model, exocytosis is targeted by a linker or adapter-based mechanism, but with targeted vesicles of a wide range in size. A combination small and large vesicles that are captured by microtubule-attached linkers might yield a flatter distribution of membrane-to-membrane contact distances and thus locations of proteins mediating membrane fusion. Our previously reported evidence that vesicles containing CSC’s^7,59^ can physically interact with cortical microtubules, together with identification of CSI-1 as a linker between CSC’s and microtubules are consistent with such a model. However, the existence of additional linker proteins mediating a wider range secretory traffic would be needed for this model to account for the observed distribution of bias for KEULE localization. Alternatively, microtubules might organize a membrane domain, which in turn could recruit upstream exocytotic proteins such as components of the exocyst complex, therefore elevating their location concentration. Hints that such microtubule-organized membrane domains may exist are suggested by observations of linear domains of protein absence that correspond to the locations of cortical microtubules, for example, by the actin polymerase Formin1, which reveals narrow “dark” linear domains defined by cortical microtubules where fluorescently-tagged Formin1 is excluded^60^. Further, EXO70, an essential exocyst protein, has been observed to localize to microtubule bundles in differentiating xylem cells, a localization shown to be dependent on the function of novel vesicle tethering proteins VETH1 and VETH2, and conserved oligomeric Golgi protein 2 (COG2)^59^. However, similar localization of EXO70 to cortical microtubules in non-xylem cells was not detected^61,62^. Several features of our quantitative measurements favor the membrane domain model over strict vesicle-microtubule tethering. Most notably, the flat probability distribution across the entire ∼180 nm enrichment zone is difficult to reconcile with direct vesicle capture at microtubule surfaces, which would be expected to produce peak enrichment at or very near the microtubule with monotonic decay. Instead, the abrupt transition from flat positive bias to negative bias at 180 nm suggests a discrete membrane domain boundary. Additionally, the substantial width of the enrichment zone (∼180 nm to each side) extends well beyond the reach of typical protein tethers, further supporting a domain-based mechanism where microtubules organize membrane composition rather than directly capturing vesicles. Applying the methods we used for KEULE offers a more powerful way to determine if exocyst or other upstream exocytotic proteins may be enriched in the microtubule-determined domains for active exocytosis revealed by our data.

One possibility for specialized exocytotic traffic targeted by cortical microtubules are cargoes involved in cell wall biosynthesis and cell wall expansion. Supporting this idea is that KEULE has been shown to be an essential partner of the Qa-SNARE SYP121, which is implicated by a growing body of evidence in mediating exocytosis required for cell growth and expansion^20,37,38,63,64^. Cellulose is synthesized at the plasma membrane, while many other structural oligosaccharides, such as pectins^65^, and wall-active proteins, such as expansins^66^, are secreted. Targeting this exocytotic traffic by cortical microtubule arrays to the same locations where newly synthesized cellulose microfibrils are also being laid down could help coordinate the assembly of these diverse components to build a functional cell wall. In addition to coordination of wall assembly, it is possible that the patterned delivery of cell wall oligosaccharides and cell wall active proteins could itself act to support the anisotropic expansion of cell walls required to create specific cell shape. Targeting the delivery of proteins and other molecules that promote wall remodeling by ordered microtubule arrays could create linear and parallel microdomains of increased wall extendibility that lie perpendicular to the main axis of the cell growth - shaping the anisotropy of cell wall expansion in the same direction as proposed in prevailing models for constraint of expansion by cellulose microfibrils organized by microtubules^14^. As discussed in the introduction of this manuscript, an important reason to consider alternative functions for cortical microtubules arrays in guiding cell wall expansion and cell shape is that experimental evidence suggests that guidance of cellulose deposition and CSC delivery alone is not sufficient to explain the anisotropy of cell wall expansion and cell growth. Microtubule targeting of cell wall-active factors by exocytosis could also help explain redirection of cell growth in response to signals, such as blue-light stimulation of phototrophic growth. Perception of blue light by Phototropin receptors causes a rapid 90° reorientation of cortical microtubule arrays^67^, but the following redirection of expansion cannot be easily explained by directed cellulose deposition model since previously synthesized cellulose is deposited perpendicular to the direction of expansion. Microtubule patterning of cell wall-active factors might allow for redirection of cell expansion regardless of cellulose orientation.

The fact that growing plant cells have a planar microtubule array that is associated with the plasma membrane, and that exocytotic events as marked by KEULE protein are of relatively low density, has provided an excellent platform to study how non-centrosomal microtubule arrays can pattern exocytosis, and do so at the level of individual exocytotic events. The templated mechanism of fine-scale exocytotic targeting revealed by our studies might be considered in other systems where there is indirect evidence that microtubules regulate exocytosis where they interact with the plasma membrane, such as in focal adhesions^1^, neurons^68–73^. and other systems with non-centrosomal cortical microtubule arrays.

## Materials and Methods

### Generation of transgenic lines

The *pKEULE-tdTomato-KEULE* construct was generated by modifying the Gateway-compatible vector pMDC43^74^. A 2049-bp region immediately upstream of the start codon of *KEULE* (At1g12360) was amplified with primers CGTTTAAACTACAGTTCCATTGGAGCAAC and TAGGTACCTGCGATCTAACAATTT CAGACTTC. The tdTomato gene was amplified from a donor vector^75^ with primers CAGGTACCATGGTGAGCAAGGGCGAGGAGGT and TGGCGCGCCCTT GTACAGCTCCTCCATGCCGT. The 35S promoter of pMDC43 was exchanged with the *KEULE* upstream regulatory region via PmeI and Acc65I sites, and GFP in pMDC43 was exchanged with *tdTomato* via KpnI and AscI sites. The full-length *KEULE* gene was amplified from genomic DNA using primers GGGGACAAGTTTGTACAAAAAAGCAG GCTTGACCATGTCGTACTCTGACTCC and GGGGACCACTTTGTACAAGAAAGCTGGG TTTATTTGGAGATCGTCTA and then inserted into the modified pMDC43 vector using Gateway cloning technology (Invitrogen) as described previously^67^. The construct was introduced into plants heterozygous for the *keule* MM125 deletion allele^40^ via Agrobacterium-mediated transformation^76,77^. T2 plants were genotyped for the *KEULE* locus using PCR: primers CGACCTTATCAGAGAGCAAGG and Keu-as0 (ACAGGGATTTGATTTGAGATCTGG) amplified a 1661-bp fragment from the wild-type allele and a 1505-bp allele from the deletion allele. Additionally, the primer pair GTCTGTTAACGGCTGCAGAA and Keu-as0 was used to amplify a fragment (1103 bp) only present in the wild-type allele. A T2 plant homozygous for the deletion allele and hemizygous for the *pKEULE-tdTomato-KEULE* transgene was isolated. A doubly homozygous line was subsequently isolated in the T3 generation and used for experiments.

To generate double-labeled plants expressing markers for KEULE protein and cellulose synthase, homozygous *pKEULE-tdTomato-KEULE/ keule* MM125 plants (ecotype Landsberg) were crossed with homozygous *pCESA6-Citrine-CESA6/cesa6prc1-1* plants (ecotype Columbia^15^. An F2 line was selected that was homozygous for both mutations and hemizygous for both markers. In the F3 generation, a quadruply homozygous plant was isolated together with a plant homozygous for both mutations, homozygous for *pCESA6-Citrine-CESA6* and hemizygous for *pKEULE-tdTomato-KEULE*. Seed from the latter plant thus show segregation for the *keule* mutant phenotype.

To generate double-labeled plants expressing markers for KEULE protein and microtubules, *p35S- EGFP-TUA5* was introduced into *pKEULE*-*tdTomato-KEULE/keule-*MM125 plants via Agrobacterium-mediated transformation. T2 seedlings expressing the microtubule label were used for experiments.

### Growth

Seedlings were grown in the dark on vertical agar plates as described in Lindeboom et al.^67^. For hypocotyl elongation assays, seedlings were grown for 5 days. For live cell imaging experiments, seedlings were imaged 3 days after germination.

### Drug treatments

Oryzalin, taxol and isoxaben were dissolved in DMSO for 100x stock solutions. Oryzalin was used at a final concentration of 20 µM, taxol (Sigma) at 20 µM and isoxaben (Sigma) at 100 nM in plant growth media (Hoagland’s No. 2). Mannitol was dissolved in plant growth media directly at a concentration of 200 mM. For control treatment (mock), DMSO was added to the growth media to a final concentration of 0.01%. Incubation of the seedlings was performed in 6-well plates (WVR, nontreated) with 2 mL of plant growth media in each well. The plates were wrapped in aluminum foil to block light and incubated at 22 °C. The drug treatment durations were 16 hours for oryzalin, and 2 hours for all other treatments. Likewise, the mock treatments were 16 hours for comparison to oryzalin, and 2 hours for comparison to the remaining treatments.

### Confocal microscopy

Imaging was performed on an instrument featuring a CSU-X1 spinning disk head (Yokogawa Electric), a DMI6000 B inverted microscope (Leica), a 100X/1.4 NA oil immersion objective (Leica), an Evolve 512 EMCCD camera (Photometrics) and a 1.2X telescope between the spinning disk unit and camera. EGFP/CITRINE and tdTomato were excited at 488 nm and 561 nm respectively, by solid-state lasers (Coherent Cube). Switching, shuttering and intensity of these lasers was performed by direct electronic control. Emission filtering was accomplished with bandpass filters, (525/50 nm for EGFP and CITRINE and 636/37 nm for tdTomato (SemRock), together with a 405/488/568 nm dichroic filter (SemRock). Exposure times were 400 ms for tdTomato-KEULE, 200 ms for EGFP-TUA5 and 800 ms for Citrine-CESA6. Images were acquired at 1 second intervals in experiments in which both tdTomato- KEULE and EGFP-TUA5 were imaged and 2 second intervals in experiments featuring tdTomato- KEULE together with Citrine-CESA6. All light energies were approximately 4 mW as measured at the end of the optical fiber entering the confocal head. For high-speed acquisitions of tdTomato-KEULE, image streaming was performed at an acquisition rate of 5 Hz (200 ms exposures). The focal plane in all experiments was stabilized by hardware Adaptive Focus Control (Leica) to mitigate changes in specimen intensity over time caused by thermal and mechanical drift in focus.

Photobleaching was performed by a Vector laser scanning unit (Intelligent Imaging Innovations) integrated into the setup above at the right camera port. Focused laser light was pulsed and scanned (pulse duration = 10 ms, raster block size = 20 units) on the image plane by a pair of galvanometer-driven mirrors inside the Vector unit. Scanner positions were calibrated to image coordinates using an automated procedure in Slidebook software (Intelligent Imaging Innovations). Citrine photobleaching was performed using the 488 nm laser at the same power as used for imaging but focused on the image plane. Areas of ∼200 µm^2^ were bleached in 20-30 s.

All imaging was performed on the upper hypocotyl cells (1-2 mm below the apical hook). Specimen mounting was performed as described previously^67^.

### Image registration

To correct for lateral specimen drift in the movies, we performed image registration in MATLAB. We first performed a running average on the image sequence using a three-frame window (yielding an image series 2 frames shorter than the original series). We then performed an intensity-based image registration algorithm using the MATLAB function imregtform, using tformtype “translation” and conserving the pixel size. To minimize interpolation artifacts, we first determined the transformation matrix that yielded the best intensity match for each consecutive image pair, then calculated the cumulative transformation matrix for the whole averaged image series and applied it to the original movie that was not frame averaged, minus the first and last frame that were lost because of the frame averaging. Some of the KEULE- tdTomato and Citrine-CESA colocalization movies (3 out of 8) still exhibited sample drift after intensity-based registration. In those cases, we applied a feature-based method employing the locations of detected KEULE spots (See Methods: Particle Tracking for details) as registration features and using the MATLAB function estgeotform2d and a rigid transform.

### KEULE particle tracking, streaming experiments, image series with labeled microtubules

Particle detection and tracking was performed on the registered movies. For streaming experiments and experiments featuring labeled microtubules (1 second time intervals) we used µTrack^49^ to detect and estimate the positions of tdTomato-KEULE clusters in each frame and subsequently link detected foci from frame to frame to track individual KEULE clusters over time. For each experiment type, we used the following parameters:

#### Streaming image series

For detection: initial psfSigma = 1.2 pixels, alpha for local max 0.05 and 10 max iterations for psfSigma estimation.

For tracking: max detection gap = 2 frames, min track length = 4 frames, max search radius = 2 pixels, gap penalty = 4.

#### 1-second interval image series

For detection: initial psfSigma = 1.2 pixels, alpha for local max 0.05 and 10 max iterations for psfSigma estimation.

For tracking: max detection gap = 1 frame, min track length = 1 frames, max search radius = 3 pixels, gap penalty = 4.

### KEULE and CESA particle tracking in dual labeled movies

In experiments assessing KEULE localization and dynamics with respect to cellulose synthase complex (CSC) trafficking and we used the Spots module in Imaris software (Bitplane) to detect and track both tdTomato-KEULE clusters and Citrine-CESA6 particles. The Imaris Spots detector and filters performed better for Citrine-CESA6 signals, in particular. CSCs as labeled by Citrine-CESA6 were detected using a size value of 0.3 µm and quality value of 110. Tracking was performed by applying the Autoregressive Motion algorithm (maximum distance = 0.35 µm, maximum gap size = 2 frames). Successful deliveries of CSC complexes to the plasma membrane were determined by dynamic criteria as described previously^7^. Briefly, a delivery event was defined when puncta that were mobile in the streaming cytosol (1) stabilized positionally in the optical plane of the plasma membrane, and (2) showed a transition to slow, steady and linear movement in the velocity range (∼200-450 nm/min) we previously have determined be characteristic of CSC’s actively synthesizing and extruding cellulose fibers into the cell wall^15^. To make this assessment, we first filtered CSC tracks by duration (>30 seconds), then used a custom GUI in MATLAB to manually inspect and verify each Spots track and associated image data for these characteristics. As a part of this process, the frame in which a tracked particle first showed evidence of stabilized lateral position was marked as the “tethering” frame – the frame showing the time and location of first stable interaction with the cell cortex of a presumed vesicle carrying one or more labeled CSC’s.

Clusters of tdTomato-KEULE were detected with Spots module using a size parameter value of 0.3 µm, a quality value of 100 and were tracked using the Connected Components algorithm (frame to frame displacements of KEULE signal are small and not auto-correlated). These settings adequately detected and tracked KEULE clusters based on visual inspection of the results.

### Maximum Likelihood Gaussian fitting for particles

We employed maximum likelihood Gaussian fitting as described in Mortensen et al.^50^ to use photon counts in estimating the accuracy of particle position and shape. To determine the number of photons collected in each image pixel, we first needed to calibrate our camera gain (photoelectrons per grey value or ADU). Photon detection can be modeled as a Poisson process, which creates a distribution in which the mean equals the variance. Therefore, the camera gain is equal to the coefficient between the mean and variance of the gray values in the image. To calculate this coefficient, we first measured the noise floor of the camera by taking the average of the gray values (intensity) of images acquired with the camera capped. We then determined the grey value variance by subtracting one image from the other then calculating the variance among these pixel-by-pixel difference values. To obtain the final variance, we divided this number by 2 to account for the fact we are using two images for determining the differences. Finally, we calculated the mean intensity of the images, subtracted the noise floor, and divided by the variance to estimate the camera gain.

For each time point in every track, we cropped out a pixel grid with the estimated center of the detected KEULE spot in the middle pixel (9 x 9 pixels). Along with a cropped image, the fitting routine requires a set of initial parameters: xCoordinate, yCoordinate, point spread function standard deviation (PSF SD), background, and an estimate of number of photons that comprise the imaged PSF. The x- and y-coordinates we used are the estimates of position from the particle tracking described above. Based on the approximate emission wavelength and objective numerical aperture, we used 133 nm for the initial standard deviation of the PSF. We estimated the background by calculating the median intensity of the outer border of the pixel grid used for fitting after subtraction of the camera noise floor. If the center of the point source is in the middle of the pixel and the PSF SD is 133 nm, we expect 0.16 of the photoelectrons to be in the center pixel. Therefore, we estimated the number of photoelectrons in the imaged spot by first subtracting the noise floor and pixel background from the intensity in the center pixel and multiplying the result by 1/0.16 (6.25). Before performing fitting using these initial data, we needed to address noise that could have been introduced by bilinear interpolation during the image registration that was employed to mitigate specimen drift. To accomplish this, we used the transformation matrix from registration to map the locations of the tracked particles back to the locations in the original image sequence, we then performed MLE fitting on the non-interpolated images by employing the method of Mortenson et al.^50^ in MATLAB.

Since we used free fit parameters and there is noise in the data from photon counting, free fluorescently tagged proteins, high local particle densities and unresolved distributions of labeled structures, we excluded distance measurement from fits that were outside the expected parameter ranges. In the case of the SD of the 2D Gaussian, we found that there was a distinct population where the SD fit parameter was < 50 nm. Since our pixels correspond to 133 nm, in these fits all the photoelectrons are contained in one pixel, which is not consistent for a diffraction-limited spot at the magnification onto our camera (roughly 4 pixels). Similarly, we found that fit SD values above 2x the estimated PSF SD in most cases represented a fit containing more than one particle. Therefore, SD fit results with SD values outside 50 to 266 nm were excluded from position measurements. Fit results with estimated numbers of photoelectrons > 1000 were also considered outliers and excluded. Finally, high local particle densities the distance in some cases caused the MLEwG fit to center on a nearby particle and not the particle marked by the starting position. We therefore also did not include position information for fits where the distance between the original estimate and the MLEwG position differed more than 2 x 133 nm. See Fig. S6 for fit parameters for the tdTomato-KEULE clusters and CIT-CESA6 labeled CSCs.

### KEULE to microtubule distance measurements

To determine the position of each KEULE spot to assess the spatial relationship of KEULE accumulation with the microtubule cytoskeleton we first determined the single frame with the highest number of measured photoelectrons for each KEULE track. For tracks that were longer than 4 frames we determined the max number of photoelectrons using a running three frames average, for shorter tracks we did not use frame averaging. The rational for this selection was two-fold. First, we reasoned that maximum signal might best be associated with the site of KEULE function, with initial signal rise being indicative of KEULE recruitment and/or assembly into a membrane fusion structure, and signal loss indicating KEULE disassociation or disassembly of the fusion machinery. Second, the precision of estimated position is proportional to photon sampling. It is also relevant to note that frame-to-frame movement of KEULE tracks were small (Fig. S7).

To locate positions of microtubule signal relative to each KEULE track location, we used a rotating linear filter combined with analysis of signal peaks and a two-step fitting procedure. At the coordinates of KEULE signal in each max value frame we drew a line 21 pixels long and 1 pixel wide extending 10.5 pixels to each side of the KEULE position and repeated this process in increments of 9 degrees to cover 360 degrees (Fig. 4C). We then projected this array of lines on the microtubule channel and extracted the intensities (in photons) along each line at 1-pixel (133 nm) intervals by bilinear interpolation. To locate potential positions of microtubule signal along each sampling angle we used the MATLAB function Findpeaks with a MinPeakProminence of 25 to identify signal peaks. The 10.5-pixel distance is typically long enough to contain the peak intensity of the nearest microtubule. In fact, in many cases this length intersects multiple microtubules, but it is also possible for the end of the lines to partially intersect a microtubule with a peak outside the 10.5 pixels. For subsequent fitting steps, we included pixels from the center until the furthest peak on each side plus 3 pixels to catch the shoulder of these peaks, and at the same time, exclude signal from microtubules whose peaks are just outside the 10.5 pixel range from influencing the fitting. We then estimated the center position of each signal peak by modifying the 2D MLEwG^50^ to a quasi-1D version, modeling the microtubule as a Gaussian wall^78^ that is one pixel wide, as shown in formula 1 below.

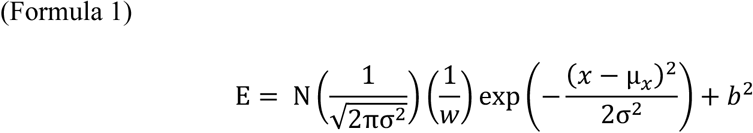

Where:

E = expected number of photoelectrons per pixel

N = total number of photoelectrons

σ = Standard deviation

𝑤 = line width in nm

𝑥 = position on the line

µ_𝑥_ = center position

𝑏^2^ = background photoelectrons per pixel

We found up to four peaks on a single line and fitted all peaks simultaneously by adding an additional term describing each additional peak. Formula 2 describes the case for fitting the case for two peaks:

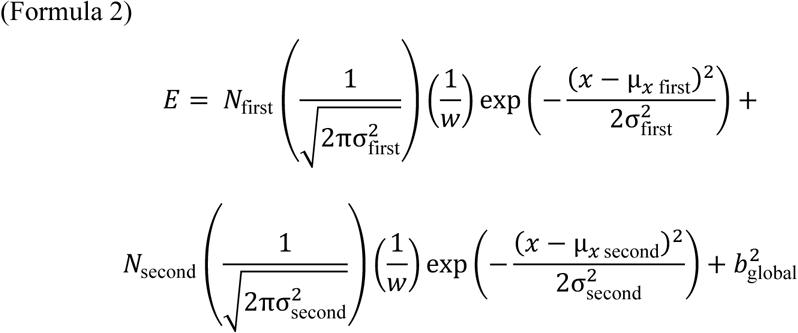

For estimating the seeding parameters for the fit, we used the center of the maximum value pixel for each peak identified by Findpeaks. We used 160 nm (median value of the peak population) as the estimate for the standard deviation and we approximated the background by taking the square root of the lowest 10 percentile of all the pixels for all angles after subtracting the noise floor.

After fitting all microtubule peaks along all angular steps, we determined the distance from the KEULE location to the center of the nearest microtubule signal. We did this in three steps. First, we determined the nearest microtubule signal peak to the KEULE location from among the initial angular samples. Second, we determined the angle of the microtubule identified by the closest peak. Measuring the distance from the KEULE location to the center of the microtubule signal is most accurate along a line perpendicular from the KEULE location to the microtubule. By determining the local microtubule signal angle, we could then determine the optimal perpendicular sample angle. To find the local orientation of the microtubule we rotated a line of 21 pixels long around the center of the location where we initially found the closest microtubule peak. We extracted the summed signal intensities along these lines and determined the angle at which we found the maximum intensity, corresponding to the microtubule angle^69^. Finally, once we established the microtubule angle, we used the angle perpendicular to that microtubule angle to extract the nearest microtubule fit for the representative KEULE position. Distance values where the fit parameters for the microtubule positions were the number of photoelectrons were < 50 or > 10000, as well as SD fit parameters < 50 or > 250 nm were excluded from distance measurements. We required all the microtubule fits on the line to be within these parameters to be included in the analysis. See Fig. S8A-C for fit parameters for KEULE (A) and microtubule signal (B) fits after selection as well as the standard error of the distance measurement between them (C).

### Analysis of KEULE localization with CESA6 delivery sites

KEULE and CESA6 spots were detected and tracked in Imaris as described above and maximum likelihood 2D Gaussian fitting was applied to each set of tracks. CESA tracks tend to be relatively long and can be confounded by a dynamic background produced by highly motile CESA containing organelles including Golgi bodies and Trans-Golgi network compartments. This background makes fully automated detection and quantitation of CSC delivery events challenging. We therefore designed a graphical user interface in MATLAB that allowed us to efficiently curate potential CESA delivery events identified in Imaris and verify whether individual tracks were indeed depicting a CESA delivery. We have shown previously that CESA delivery to the plasma membrane typically shows three distinct phases; (1) erratic movement of CESA containing vesicles as they appear in the optical plane at the cell cortex, (2) positional-stabilization and (3) slow and steady movement with typically linear trajectories. We speculate that the second phase represents molecular tethering at the plasma membrane and the events that lead to vesicle fusion and cargo delivery. The third phase is behavior that we have shown is characteristic of active CSC complexes in the plasma membrane as they synthesize cellulose^15^. Thus, observation of this latter behavior indicates successful delivery of CESA cargo to the plasma membrane. For each CESA track that showed these three distinct phases, we marked the first frame of CESA stabilization at the plasma membrane. From this position, we determined the distance to the nearest KEULE spot in a 30- frame time range (60 seconds). Some degree of coincidence between CESA deliveries and KEULE spots is expected by chance alone, especially since the density of KEULE clusters at the plasma membrane is relatively high. To evaluate whether the observed distribution of KEULE locations with sites of CESA stabilization might be due to chance alone, we generated random positions in the same regions of interest used for the experimental data and calculated the distances to the nearest KEULE spot. We used these simulated data to model the expected distribution of distances and compared this distribution to the distribution of distances found for confirmed CESA delivery events. First, we divided both the experimental and simulated distance distributions up into 40 nm distance bins and calculated the bin-by-bin count frequencies by the total number of simulated counts. We then introduced a bias parameter 𝛿 that equates an observed frequency in each distance bin to the expected value from the simulated distribution as shown in formula (4).

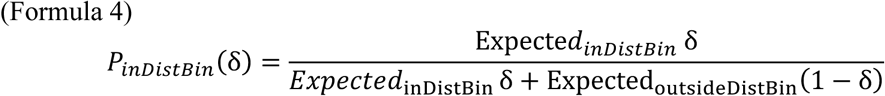

The value of 𝛿 may vary between 0 and 1. For a possible observed frequency that equals the expected frequency, 𝛿 is equal 0.5. If 𝛿 is greater than 0.5, an observed frequency is higher than expected. Likewise, if 𝛿 is less than 0.5 the observed frequency lower than expected under the null hypothesis. Use of the parameter allows us to use a Bayesian approach to test multiple hypotheses to ask what bias value best may best explain the actual observed frequencies. To do this, we calculated the Likelihood Ratio of observing an experimental result given a range of values for 𝛿 (in increments of 0.001, excluding 0) for each frequency bin:

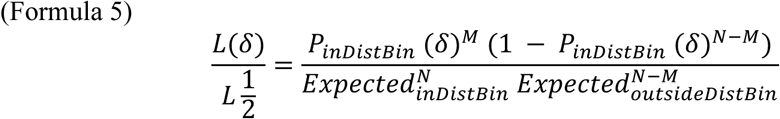

Where:

𝐿(𝛿) = likelihood of the data given the bias 𝛿,

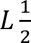= likelihood of the data in the neutral case (no positive or negative bias)

𝑁 = total number of measurements

𝑀 = number of measurements in this specific distance bin

We then found the maximum a posteriori probability (MAP) for 𝛿 and computed the shortest range of 𝛿 values for the highest posterior density (HPD) that contains 95% of the normalized probability density. Maximum likelihood estimates for 𝛿 below 0.5 indicate underrepresentation at the distance of interest and above 0.5 indicate overrepresentation of experimental data compared to the unbiased case.

### Analysis of KEULE localization with microtubules

We used the same Bayesian approach describe above for analyzing the localization of KEULE clusters with respect to the estimated center of the nearest microtubule. We changed the distance bin size from 40 to 20 nm.

### Software

We have developed and released four complementary MATLAB toolkits that together provide a pipeline for analyzing spatial relationships in fluorescence microscopy:

- MovieRegistration (https://github.com/JelmerLindeboom/MovieRegistration) for drift correction
- MLE-1D-Gaussian-Fitting (https://github.com/JelmerLindeboom/MLE-1D-Gaussian-Fitting) for distance measurements to linear targets like microtubules
- CellROI (https://github.com/JelmerLindeboom/CellROI) for spatial filtering
- BayesianColocalization (https://github.com/JelmerLindeboom/BayesianColocalization) for Bayesian analysis

### Hot spot analysis

To investigate if some KEULE events are independent of each other each or not (e.g. if “hot spots” for KEULE accumulation may exist), we examined colocalization between sequential KEULE spots in time series collected from cells expressing both labeled KEULE and tubulin proteins. Within a region of interest in each time series, we selected all KEULE tracks that were first detected between frames 2 and 31 and calculated the distance to all other KEULE tracks that started subsequently up to and including frame 301. For position estimates we used the frame in the track that showed the highest number of fitted photoelectrons. To compare the degree of colocalization of the initial KEULE tracks with future KEULE tracks to that expected from chance alone, we created a simulated initial dataset with random locations within the same region of interest and calculated the distances to all KEULE tracks in the same time range.

Since KEULE foci are positionally stable (Fig. S7) we chose a stringent distance threshold of 133 nm (1 pixel) to declare two spots or locations to be co-localized. Because of the relatively high density of KEULE spots at the plasma membrane, colocalization is expected at some point in time, therefore the time frame of colocalization analysis is also important. Indeed, we observe a large degree of colocalization in the whole dataset for both the experimental and simulated data, but with very distinct waiting time distributions (Fig. S5D). Based on the differences in the distribution of waiting times between random simulated positions and KEULE positions until the next KEULE spot arrives, we determined that a 40 second time delay window captures most biologically relevant colocalizations while excluding most random colocalizations due to KEULE density and chance.

When examining patterns of KEULE foci, we also observed that sites of previous exocytosis events were 3.4 times more likely to show consecutive exocytosis events within 60 seconds than random sites at the plasma membrane (Fig. S5A-E,G). Such “hot spots” were also significantly over-represented within 180 nm of microtubules, but only modestly so (1.25 times more likely) (Fig. S5F,G).

## Acknowledgments

We thank Michael Davidson for the kind gift of the tdTomato construct. We thank Zdeněk Lánský, Johnathan Cooper-Knock and Masayoshi Nakamura for advice and insightful comments.

## Author contributions

Conceptualization: JJL, RG, VK, DWE

Investigation: JJL, RG, VK, DWE

Methodology: JJL, RG, VK, DWE

Formal analysis: JJL, DWE

Software: JJL

Visualization: JJL, RG, DWE Funding acquisition: VK, DWE

Writing – original draft: JJL, RG, DWE

Writing – review & editing: JJL, RG, VK, DWE

## Data and materials availability

Biological material, raw data, processed data are available upon request. Matlab scripts used for processing data are available at GitHub (https://github.com/JelmerLindeboom).

## Supplemental Materials

**Fig. S1.**
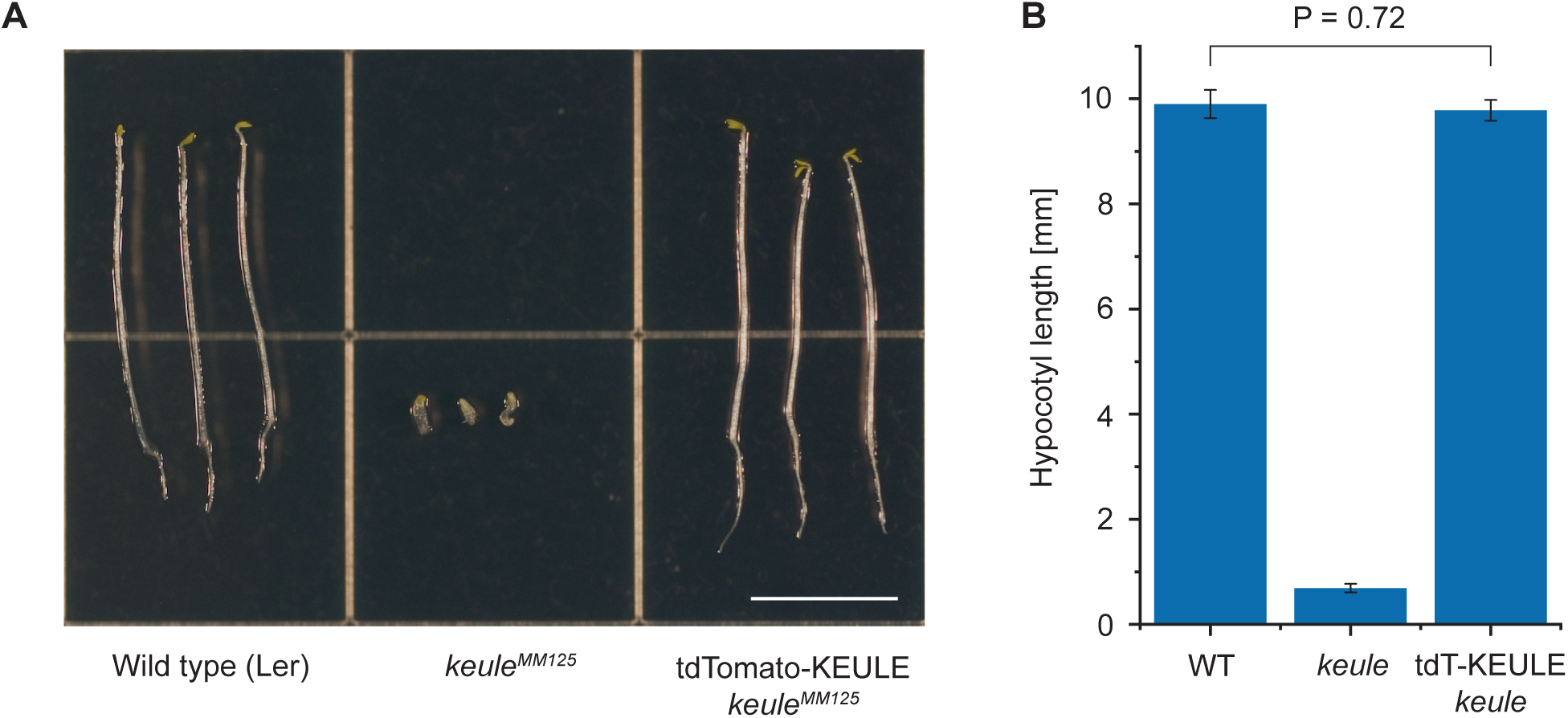
Complementation of the *keule* mutation. **(A)** Seedlings of indicated genotypes were grown in the dark for 5 d. The recessive *keule* MM125 allele is seedling lethal; mutant seedlings have bloated, multinucleate cells. Expression of a tdTomato-tagged version of KEULE, expressed under the control of the native upstream regulatory element, complemented the *keule* mutation in dark-grown seedlings. Scale bar, 5 mm. **(B)** Hypocotyl elongation in 5-day-old dark-grown seedlings. Rescued seedlings were comparable to wild-type (*n* = 50 plants per genotype, *P* = 0.72 by *t*-test). Error bars represent SEM.

**Fig. S2.**
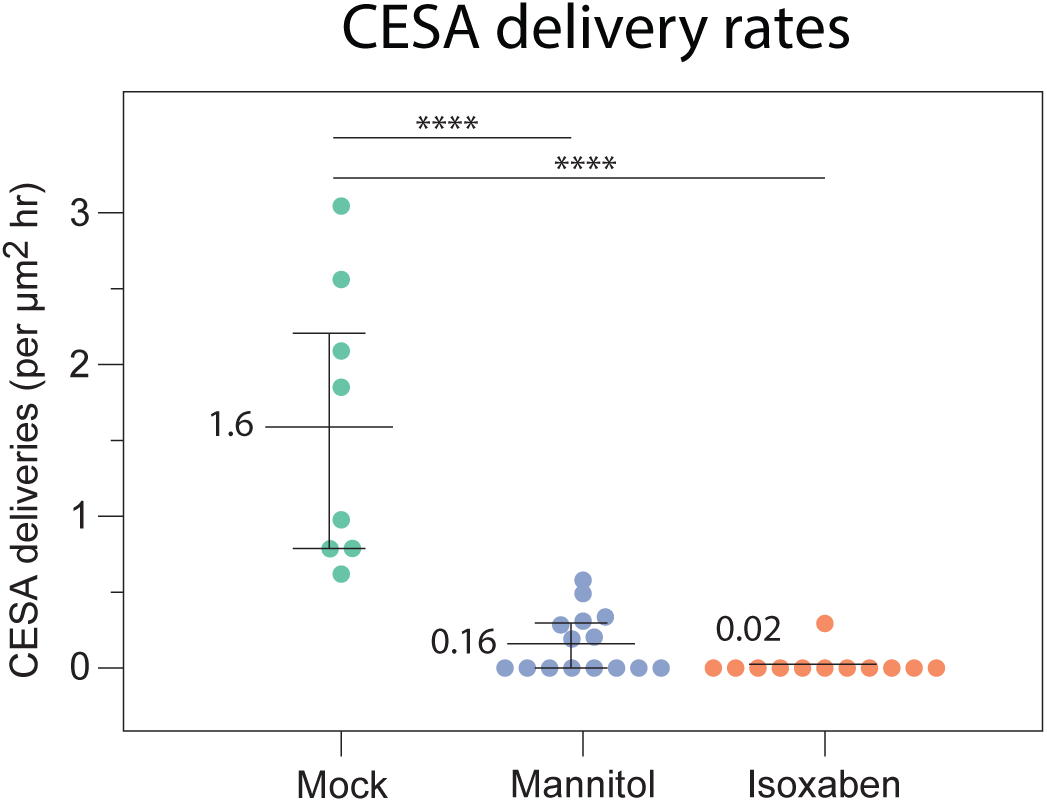
CSC delivery rates. CSC delivery rates to the plasma membrane of cells expressing tdTomato- KEULE and CIT-CESA6 after treatment with 0.01% DMSO (mock), 200 mM mannitol or 100 nM isoxaben. See Methods for treatment durations. Values are means, whiskers are quartiles. Mock: n = 8 cells from 8 seedlings, 208 delivery events observed in 1570 µm^2^ total area over 0.0828 hr. Mannitol: n = 15 cells from 15 seedlings, 11 delivery events observed in 880 µm^2^ total area over 0.0793 hr. Isoxaben: n = 12 cells from 12 seedlings, 2 delivery events observed in 996 µm^2^ total area over 0.0793 hr. **** p<0.0001, Wilcoxon two sample test.

**Fig. S3.**
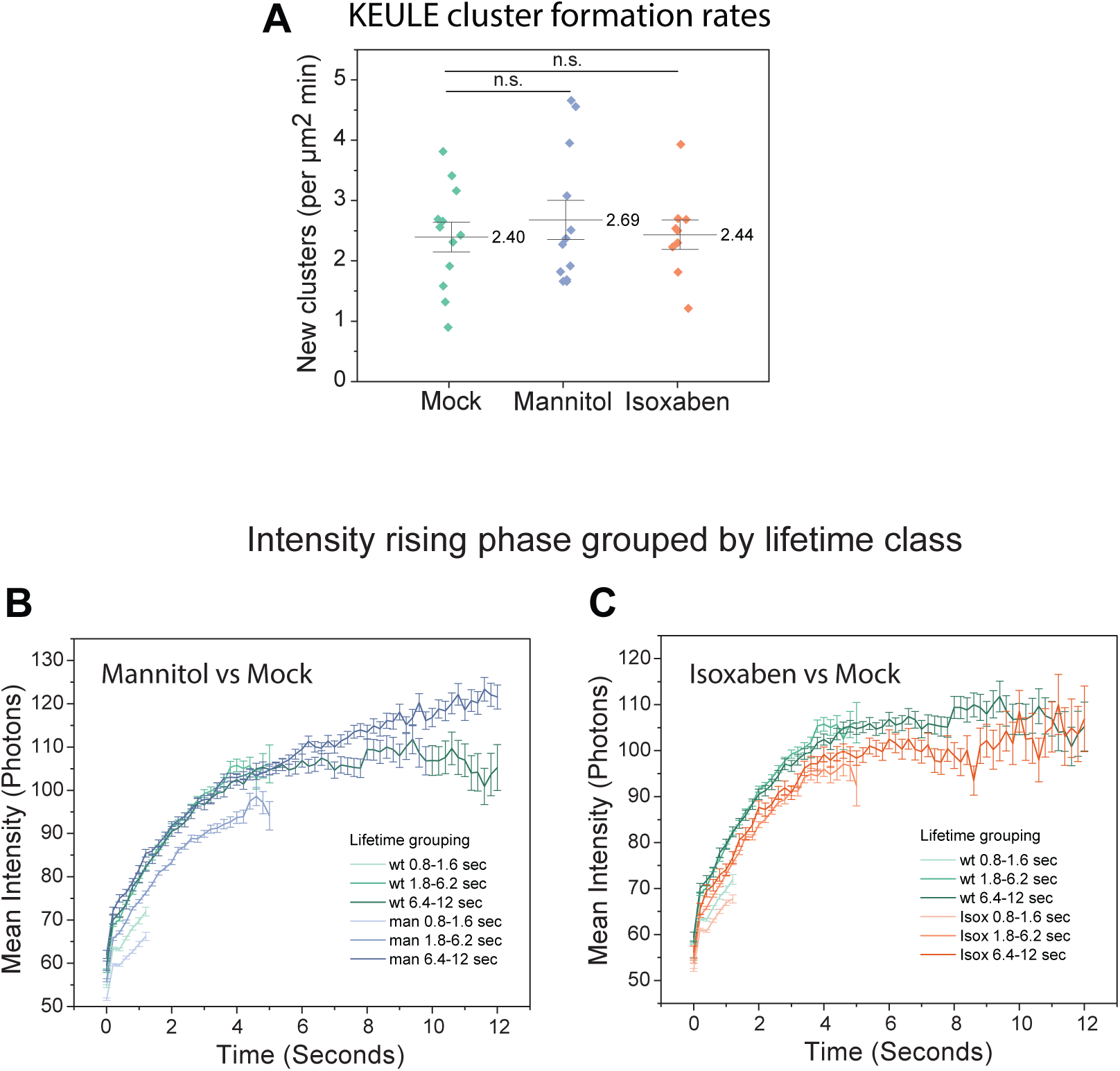
Intensity of dynamics of tdTomato-KEULE clusters by lifetime. (**A**) Average tdTomato- KEULE cluster signal intensity with lifetimes ranging from 0.8 to 6.8 s for mock, mannitol treatment. and isoxaben treatment. Whiskers are means and quartiles. n.s. p>0.05 Wilcoxon two sample test. (**B** and **C**) Comparison of average signal intensities of tdTomato-KEULE clusters in three cumulative lifetime ranges, 0.8 to 1.6 s, 1.8 to 6.2 s and 6.4 to 12.2 s, between (**B**) mock vs mannitol treatment, and (**C**) mock vs isoxaben treatment. Mock: n = 15,130 clusters observed in 12 cells from 12 plants. Mannitol: n = 35,496 clusters observed in 15 cells from 15 plants. Isoxaben: n = 25,251 clusters observed in 12 cells from 12 plants. Error bars are SEM

**Fig. S4.**
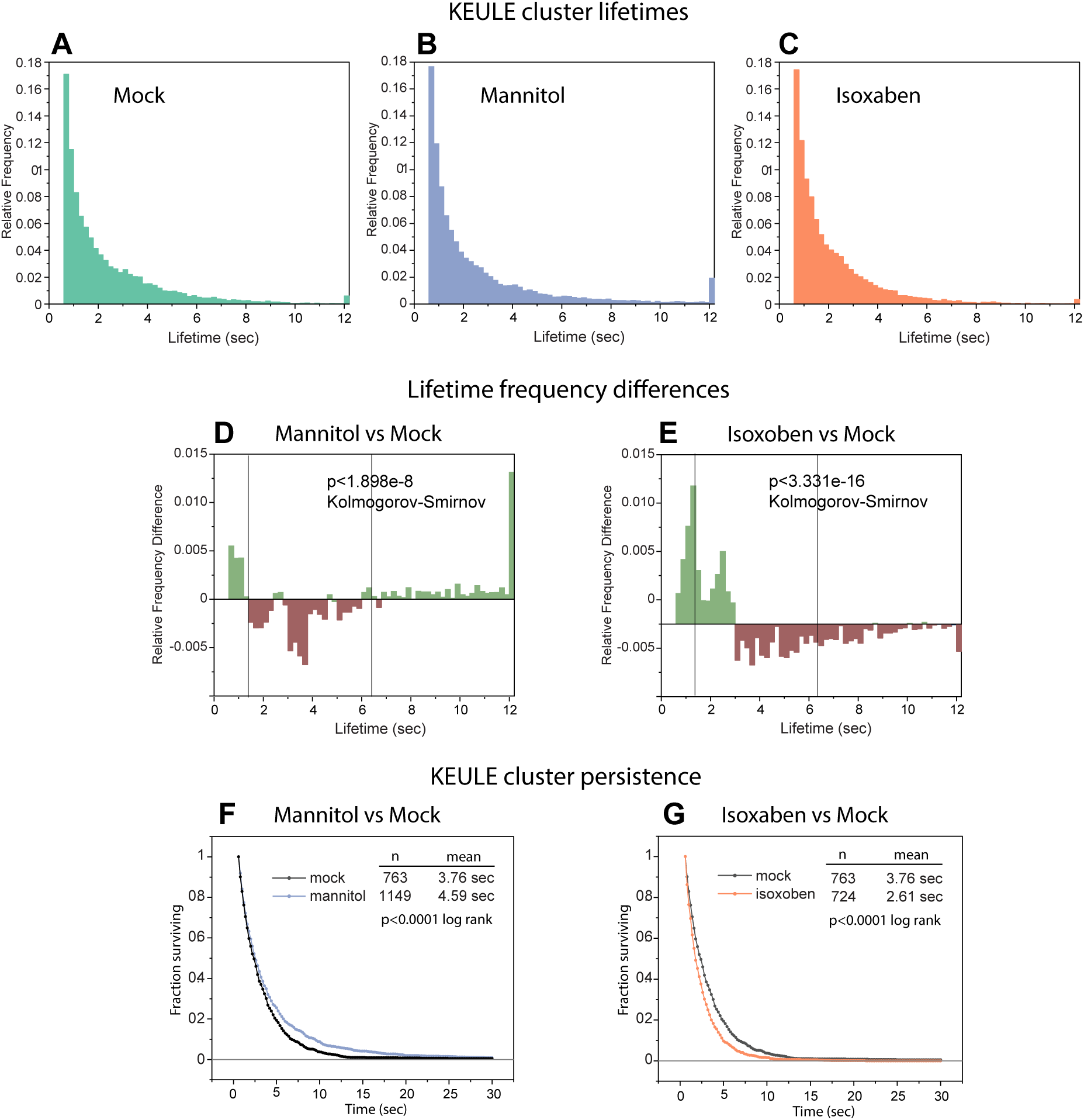
KEULE cluster lifetimes. (**A** to **C**) Lifetime distribution of tdTomato-KEULE clusters in mock (**A**), mannitol (**B**) and isoxaben (**C**) treated cells. (**D** and **E**) Difference in relative lifetime frequency between mannitol and mock (**D**), and isoxaben and mock (**E**). Vertical lines indicate the lifetime classes in Figure 1G. (**F**) Formation rates of new tdTomato-KEULE clusters. Values are means. (**G** and **H**) Comparison of the fraction of remaining tdTomato-KEULE clusters that were present in the first frame of the movie for mannitol to mock (**G**) and isoxaben to mock. Mock: n = 763 clusters observed in 12 cells from 12 plants. Mannitol: n = 1149 clusters observed in 15 cells from 15 plants. Isoxaben: n = 724 clusters observed in 12 cells from 12 plants.

**Fig. S5.**
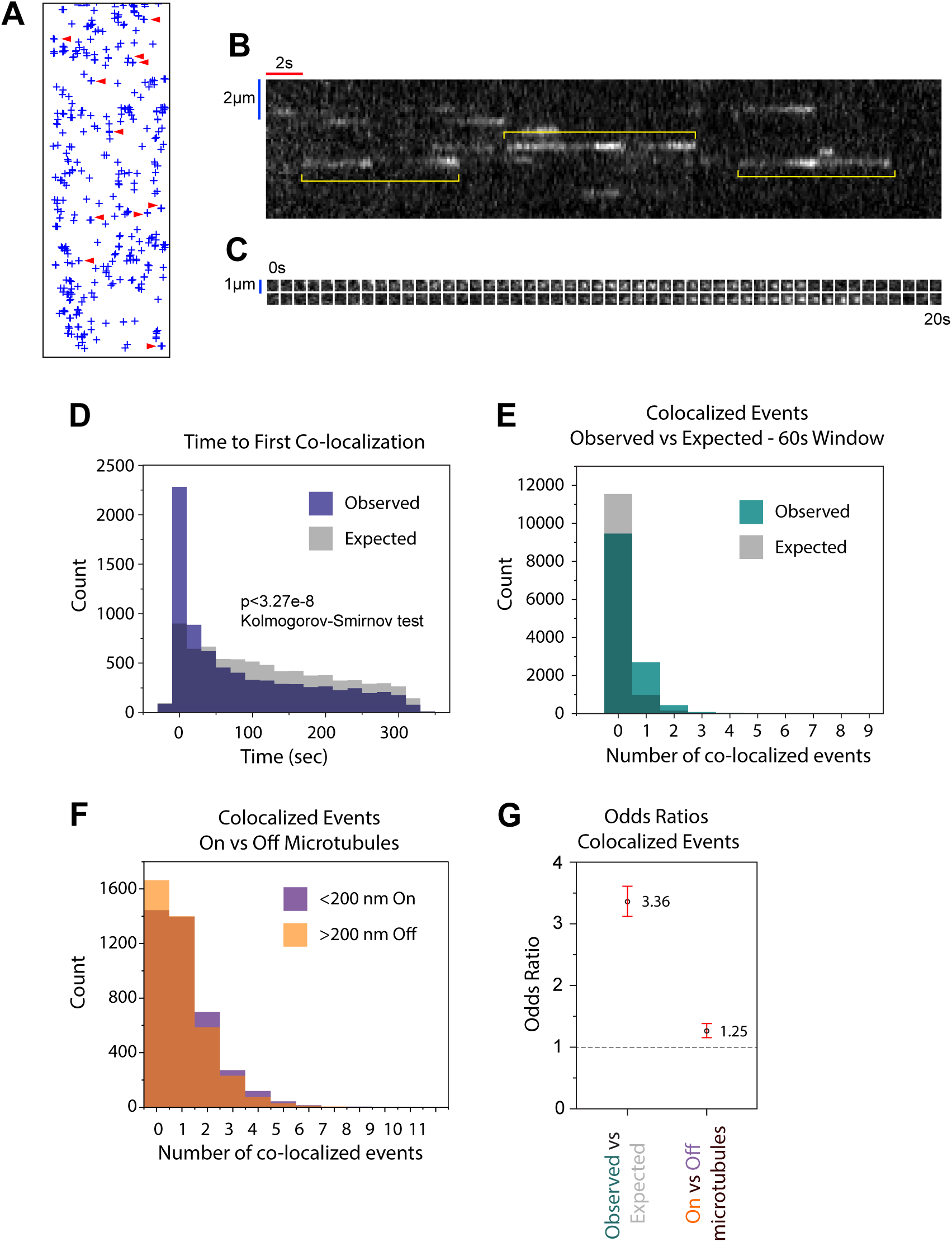
KEULE cluster hot spots. (**A**) Map of tdTomato-KEULE cluster detections over x minutes. Crosses are positions determined by MLEwG fitting. Arrowheads indicate examples of repeat detects at the same location, or hot spots. (**B**) Kymograph of tdTomato-KEULE signal along a 3 pixel wide transect line. Yellow brackets indicate examples of repeated KEULE cluster formation at the same location. (**C)** Individual image frames at a hot spot, 200 msec intervals over 20 seconds. (**D**) Distributions of waiting times for a second cluster to appear at the same location as a previous cluster, as defined by a stringent co-localization cutoff distance of 133 nanometers. The distributions for observed data and randomly simulated data are shown in purple and grey, respectively. Note that the observed intervals are significantly skewed to shorter intervals as compared to those in the simulated data (p<3.27E-8 Kolmogorov-Smirnov test). n = 12,719 clusters in 17 cells from 17 plants. (**E**) Distributions of the number of co-localized events observed at each KEULE cluster location within 60 seconds of the formation of the first cluster. Colocalization events were 3.36 more frequent in observed data (green) than expected by chance alone (grey), as determined by odds ratio analysis (see panel (**G**)). n = 9458 clusters in 17 cells from 17 plants. (F) Distributions of KEULE colocalization events either on or off microtubules on observed data. The distance threshold for microtubule association was < 200 nm. Sequential KEULE cluster co-localization was 1.25 times more frequent for KEULE events associated with microtubules than for those not associated with microtubules, as determined by odds ratio analysis (see panel (G)). n = 4000 clusters on microtubules and 4866 clusters off microtubules in 17 cells from 17 plants. (G) Odds ratios and 95% confidence intervals for the data shown in panels (G) and (F). The dotted line indicates the odds ratio expected for the null hypothesis of no different from random.

**Fig. S6.**
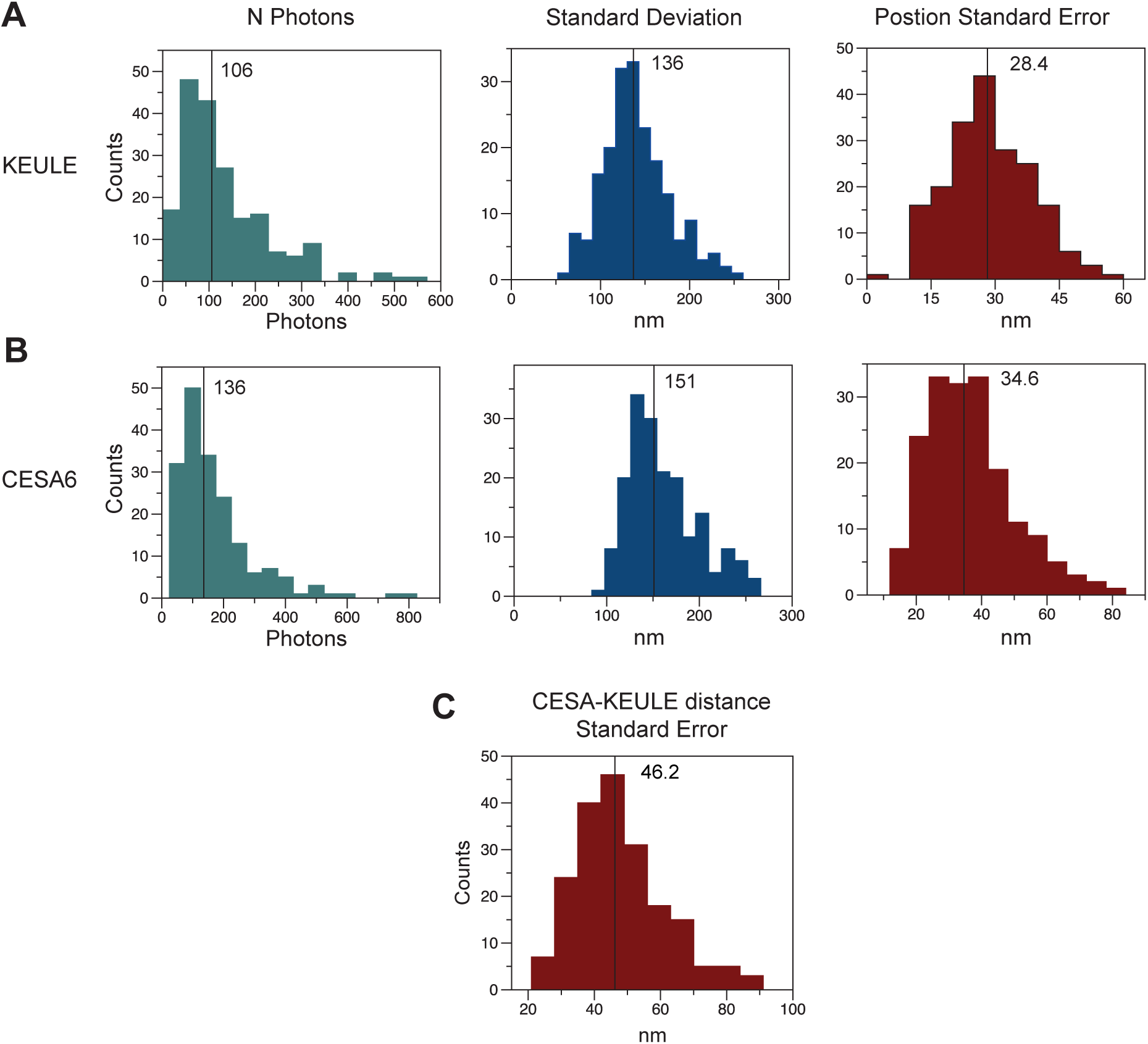
Fit parameters for CSC vesicles in the tethering frame and the nearest KEULE clusters to these tethering sites. (**A**) Distributions of fit parameters for MLEwG fits of tdTomato-KEULE clusters. (**B**) Distributions of fit parameters for MLEwG fits of CIT-CESA6 signal. (**C**) Distribution of standard errors for KEULE to microtubule distance measurements. Black lines and values indicate means. n = 194

**Fig. S7.**
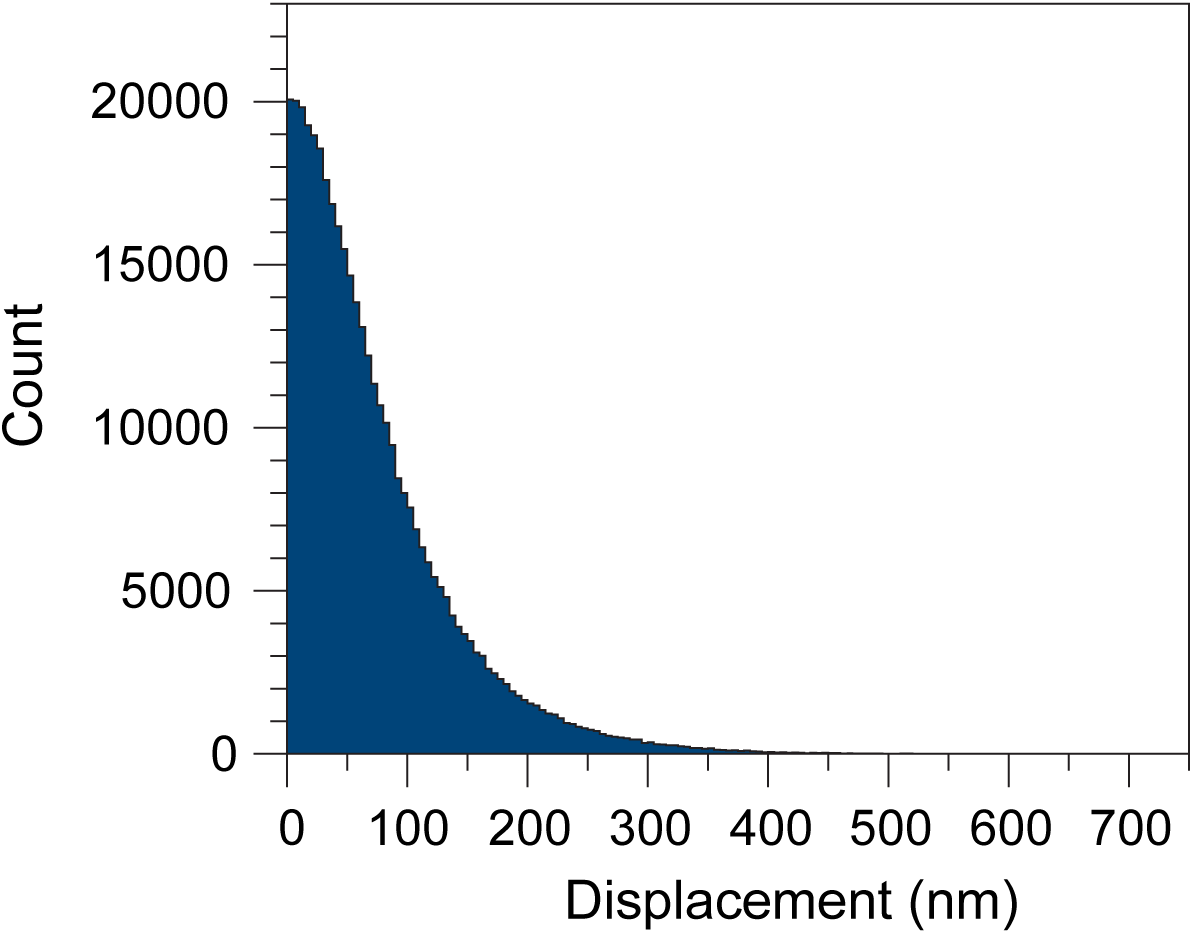
Frame-to-frame displacements of tdTomato-KEULE clusters in 200 ms streaming acquisitions. n = 283,127 displacements for 15,130 clusters observed in 12 cells from 12 plants.

**Fig. S8.**
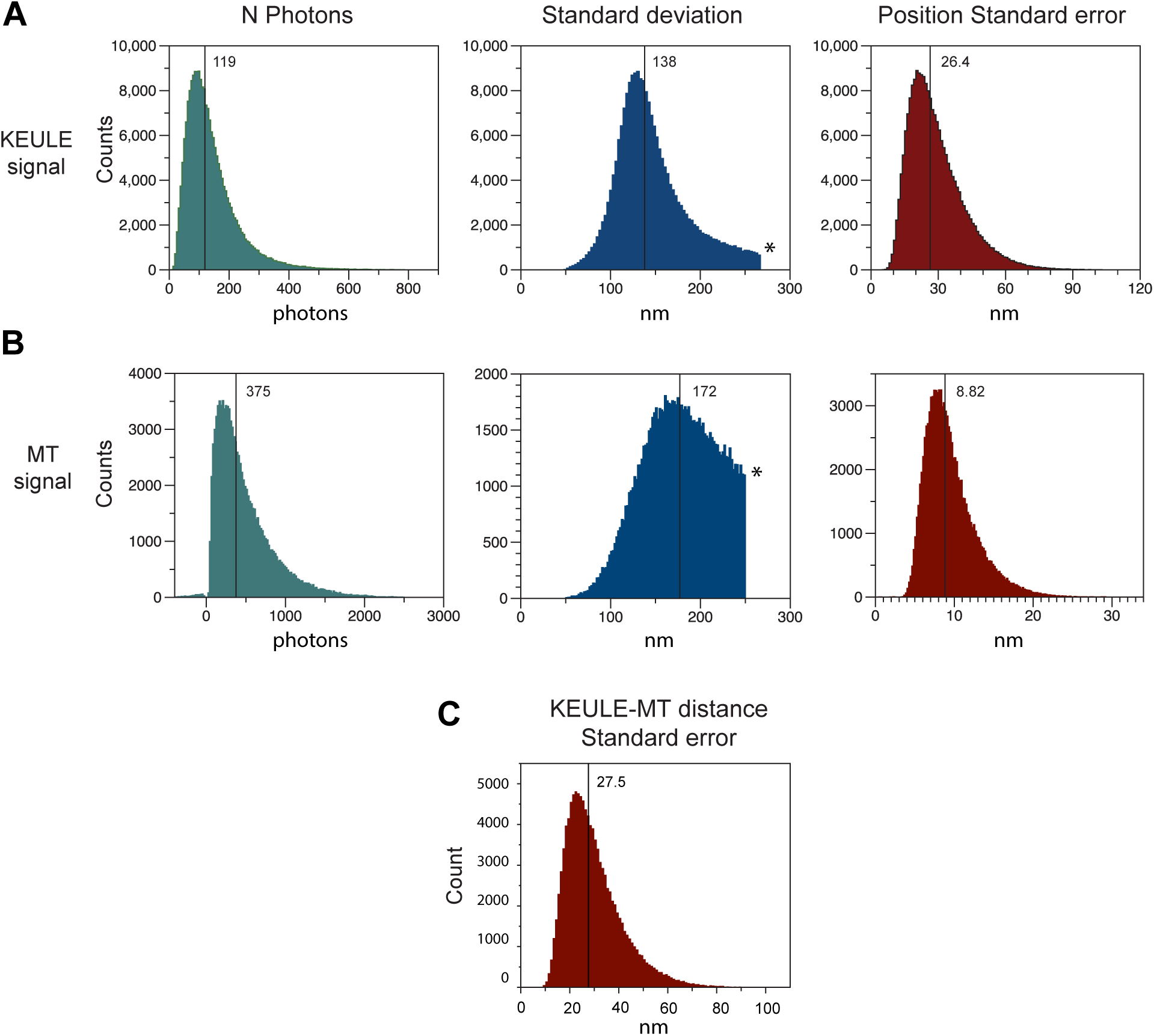
Fit parameters for KEULE to microtubule distances used for localization analysis. (**A**) Fit parameters for MLEwG fits of tdTomato-KEULE clusters. (**B**) Fit parameters for MLEw1dG fitting of microtubule signal. (**C**) Standard error of KEULE to microtubule distance measurements. Black lines and values indicate means. *Truncations of standard deviation distributions are due to cutoff values used to filter and exclude bad fits due to signal patterns that complicate localization analysis. Bad fits for tdTomato-KEULE signal occur where more than one tdTomato-KEULE cluster are close in proximity, and bad fits for microtubule signal occur where microtubules are dense and where they intersect in complex patterns. n = 115,963 KEULE clusters, nearest microtubule locations and KEULE-microtubule distances observed in 14 cells from 14 plants.

**Movie S1.**

**tdTomato-KEULE cluster dynamics.** Spinning disk confocal images of tdTomato cluster formation and loss at the plasma membrane of an epidermal hypocotyl cell in a dark grown *Arabidopsis* seedling. To reduce noise, images were averaged with a running 3-frame window. Streaming acquisition at 5 frames/sec. Playback at 4x real time.

**Movie S2.**

**MLE Gaussian fits of tdTomato-KEULE clusters.** Spinning disk confocal images of tdTomato- KEULE clusters in an epidermal hypocotyl cell of a dark grown *Arabidopsis* seedling. Magenta circles indicate an automated detection of a KEULE cluster. For each of these detects the coordinates of the signal centroid number of detected photons were estimated by MLE Gaussian fitting (see methods). Streaming acquisition at 5 frames/sec. Playback at 10x real time.

**Movie S3.**

**KEULE localization to CSC delivery events.** Spinning disk confocal images of co-expressed tdTomato- KEULE and Cit-CESA6. Filled arrowheads point to locations of tracked CSC signal. Empty arrowheads point to co-localized KEULE clusters that appear shortly after appearance and positional stabilization of CESA6 signal. 2 second acquisition intervals. 3-frame running average. Playback at 10x real time.

**Movie S4.**

**MLE Gaussian fits of tdTomato-KEULE clusters and EGFP-TUA5 labeled microtubules.** Spinning disk confocal images of co-expressed tdTomato-KEULE and EGFP-TUA5 labeled microtubules. White circles indicate automated KEULE signal detects. The position of each cluster at the first frame of stabilization was estimated at a sub-pixel level if MLE Gaussian fitting. 1 second acquisition intervals. Playback at 20x real time.

